# Overexpression of a non-muscle RBFOX2 isoform triggers cardiac conduction defects in myotonic dystrophy

**DOI:** 10.1101/649715

**Authors:** Chaitali Misra, Sushant Bangru, Feikai Lin, Kin Lam, Sara N. Koenig, Ellen R. Lubbers, Jamila Hedhli, Nathaniel P. Murphy, Darren J. Parker, Lawrence W. Dobrucki, Thomas A. Cooper, Emad Tajkhorshid, Peter J. Mohler, Auinash Kalsotra

**Affiliations:** Departments of Biochemistry, Urbana-Champaign, IL; Departments of Physics, Urbana-Champaign, IL; Departments of Bioengineering, Urbana-Champaign, IL; Carl R. Woese Institute for Genomic Biology, Urbana-Champaign, IL; Centers for Macromolecular Modeling, Bioinformatics, and Experimental Molecular Imaging at Beckman Institute for Advanced Science and Technology, Urbana-Champaign, IL; Cancer Center@Illinois, University of Illinois, Urbana-Champaign, IL; Department of Physiology and Cell Biology, Davis Heart and Lung Research Institute, College of Medicine, Wexner Medical Center, The Ohio State University, OH; Department of Pathology and Immunology, Baylor College of Medicine, Houston, TX

**Keywords:** Myotonic dystrophy, alternative splicing, cardiac arrhythmias, genome editing, Protein-RNA interactions, microRNA, genomics, molecular dynamics

## Abstract

Myotonic dystrophy type 1 (DM1) is a multisystemic genetic disorder caused by a CTG trinucleotide repeat expansion in the 3′ untranslated region of *DMPK* gene. Heart dysfunctions occur in nearly 80% of DM1 patients and are the second leading cause of DM1-related deaths. Despite these figures, the mechanisms underlying cardiac-based DM1 phenotypes are unknown. Herein, we report that upregulation of a non-muscle splice isoform of RNA binding protein RBFOX2 in DM1 heart tissue—due to altered splicing factor and microRNA activities—induces cardiac conduction defects in DM1 individuals. Mice engineered to express the non-muscle RBFOX2 isoform in heart via tetracycline-inducible transgenesis, or CRISPR/Cas9-mediated genome editing, reproduced DM1-related cardiac-conduction delay and spontaneous episodes of arrhythmia. Further, by integrating RNA binding with cardiac transcriptome datasets from both DM1 patients and mice expressing the non-muscle RBFOX2 isoform, we identified RBFOX2-driven splicing defects in the voltage-gated sodium and potassium channels, which can alter their electrophysiological properties. Thus, our results uncover a *trans*-dominant role for an aberrantly expressed RBFOX2 isoform in DM1 cardiac pathogenesis.

## INTRODUCTION

DM1 is an autosomal-dominant disorder and the most commonly inherited form of adult onset muscular dystrophy^1, 2^. The disease arises due to an expansion of trinucleotide CTG repeat in the 3’-UTR of *DMPK* gene^3–5^, which produces mutant RNAs with long tracts of CUG repeats r(CUG)^exp^ forming stable hairpin loops that aggregate in the nucleus^6–8^. Although *DMPK* is broadly expressed, and the disease affects multiple tissues, the predominant causes of mortality are muscle wasting and sudden cardiac death^9–11^. More than two-thirds of DM1 individuals experience severe heart dysfunctions including cardiac-conduction delay, fatal sinoatrial and atrioventricular blocks, atrial fibrillation, and ventricular arrhythmias^9, 12, 13^. Affected individuals often display atrophy of the conduction system as well as fibrofatty infiltration of the myocardium and the Bundle of His. Further, some patients develop dilated cardiomyopathy along with systolic/diastolic dysfunctions^11, 14, 15^.

The two major mechanisms by which r(CUG)^exp^ RNAs cause toxicity and thereby disease pathogenesis are: (i) the accumulated r(CUG)^exp^ RNAs sequester muscleblind-like proteins (MBNL1, MBNL2 and MBNL3) with high affinity, resulting in their depletion and loss-of-function^16–18^; (ii) r(CUG)^exp^ RNAs activate the protein kinase C pathway and suppress the expression of specific micro (mi)RNAs causing upregulation of CELF1 protein^19–21^. MBNL and CELF family of RNA binding proteins direct a large number of developmentally-regulated alternative splicing and polyadenylation decisions^22–26^. Therefore, disruption of their normal activities in DM1 shifts RNA processing of target pre-mRNAs towards fetal patterns^27–31^, many of which induce key features of the disease^32–35^. The exact molecular basis for the electrophysiological and cardiac contractility abnormalities in DM1, however, is still undetermined.

Here, we demonstrate that the non-muscle splice isoform of RNA-binding protein FOX2 (RBFOX2_40_) is overexpressed in DM1 human heart tissues, and that this overexpression results from a combination of elevated CELF1 and reduced miRNA activities. Modeling the increase in non-muscle RBFOX2_40_ isoform in mouse hearts was sufficient to trigger DM1-related cardiac conduction defects. By integrating RBFOX2-RNA interactions with cardiac transcriptome datasets—from DM1 patients, CELF1 overexpressing mice, MBNL1 knockouts, and mice expressing the non-muscle RBFOX2_40_ isoform—we identified a unique set of RBFOX2_40_-driven mRNA splicing defects that are dysregulated in the DM1 hearts. Specifically, the RBFOX2_40_ isoform caused missplicing of voltage-gated sodium and potassium channel transcripts directing generation of pro-arrhythmic variants that elicit altered rates of ion transport and electrophysiological channel properties. Thus, upregulation of non-muscle RBFOX2_40_ isoform augments the production of pathogenic ion channel splice variants that may directly contribute to DM1-related cardiac conduction delay and arrhythmogenesis.

## RESULTS

### Selective upregulation of the non-muscle RBFOX2_40_ isoform in DM1 heart tissue

Considered as master regulators of tissue-specific alternative splicing^36, 37^, RBFOX proteins (RBFOX1, RBFOX2, and RBFOX3) utilize a highly conserved RRM domain to bind (U)GCAUG motifs in pre-mRNAs and control splicing in a position-dependent manner^38–40^. Mutations in *RBFOX2* gene were identified as a major risk for congenital heart disease^41–43^; and essential roles for RBFOX2 were recently demonstrated in pathogenesis of stress-induced hypertrophy, heart failure, and cardiac pathology of diabetes^44, 45^. We have determined that RBFOX2 protein levels are significantly increased in the autopsied heart samples of DM1 patients but not of individuals with a history of arrhythmias caused due to left ventricular tachycardia **(Fig. 1A, B)**. Parallel examination of *RBFOX2* mRNA unexpectedly showed only a modest increase in abundance in DM1 heart tissues **(Supplementary Fig. 1A)**.

**Figure 1.**
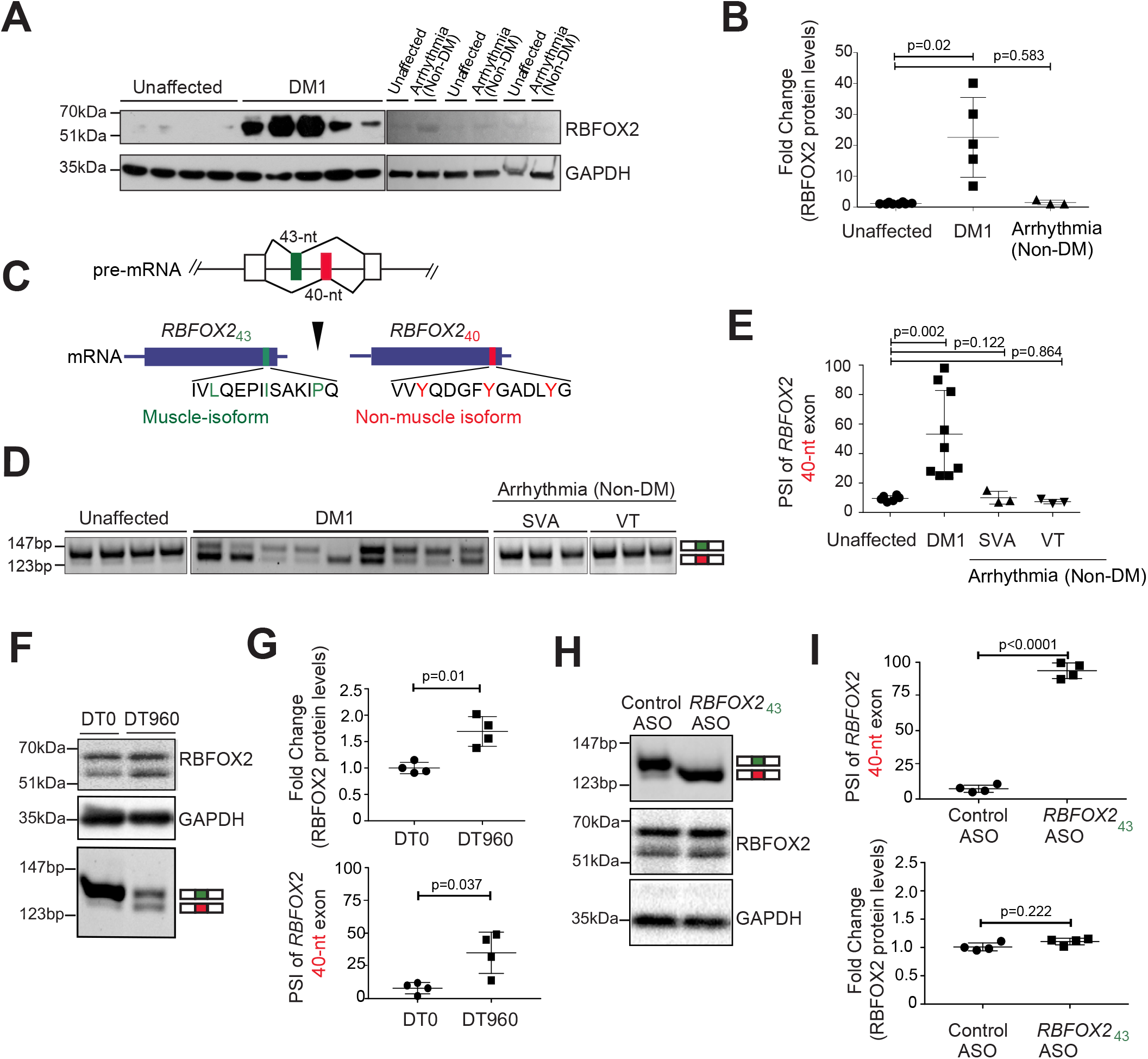
Selective upregulation of the non-muscle RBFOX2_40_ isoform in DM1 heart tissue. **(A)** Immunoblot analysis of RBFOX2 protein in unaffected (n=8), DM1 (n=5), and arrhythmic non-DM (n=3) human heart samples. **(B)** Quantification of relative band intensities for RBFOX2 from **A**, normalized to GAPDH levels. **(C)** Schematic of amino acid residues encoded by *RBFOX2* non-muscle (40-nt) and muscle-specific (43-nt) exons. Three tyrosine (Y) residues specific to the 40-nt exon are highlighted in red. **(D)** RT-PCR analysis monitoring the inclusion of *RBFOX2* 40-nt exon in unaffected (n=8), DM1 (n=9), and arrhythmic non-DM [SVA: Sustained Ventricular Arrhythmia (n=3); and VT: ventricular Tachycardia (n=3)] human heart samples. **(E)** Percent Spliced In (PSI) values of *Rbfox2* 40-nt exon from **D**. **(F)** Immunoblot analysis of RBFOX2 protein (top) and RT-PCR analysis of *Rbfox2* 43-nt and 40-nt exons (bottom) in HL-1 cells transfected with DT0 or DT960 plasmids for 48h. **(G)** Relative quantification of immunoblot and RT-PCR data from **F.** n=4 independent transfections. **(H)** RT-PCR analysis of *Rbfox2* 43-nt and 40-nt exons (top), and immunoblot analysis of RBFOX2 protein (bottom) from HL-1 cells treated with control ASO or with an ASO targeting the 5’ss of *Rbfox2* 43-nt exon for 48h. **(I)** Relative quantification of RT-PCR and immunoblot data from **H**. n=4 independent transfections. All data are mean ± s.d., and p-values were derived from parametric t-test (two-sided, unpaired), with Welch’s correction.

*RBFOX2* gene has a complex architecture comprising multiple promoters and alternative exons **(Supplementary Fig. 1B)**, which direct tissue-specific expression of various RBFOX2 isoforms that are functionally distinct in their RNA binding activities, subcellular localization, and interactions with regulatory co-factors^36^. Amongst them, a pair of 43-nucleotide (nt) and 40-nt exons in the c-terminal domain encode the muscle and non-muscle isoforms that are expressed in a mutually exclusive, developmentally regulated, and evolutionarily conserved manner. Interestingly, the 40-nt exon contains three tyrosine residues **(Fig. 1C)** that promote a higher-order RBFOX2 complex of proteins called the Large assembly of Splicing Regulators (LASR) complex, which enhances RBFOX2’s splicing activity^46, 47^. The 43-nt exon lacks these tyrosine residues **(Fig. 1C)**, and therefore, fails to form the higher-order LASR complex^47^. Both humans and mice predominantly express the 40-nt exon containing RBFOX2 isoform in fetal hearts, which is replaced by the 43-nt exon containing isoform in adult hearts, specifically within cardiomyocytes **(Supplementary Fig. 1C-E)**. Strikingly, we noticed a significant splicing shift—from 43-nt to 40-nt exon—within *RBFOX2* transcripts in DM1 patient heart samples (ΔPSI ranging from 12% to 95%), which was not detected in the hearts of individuals diagnosed with non-DM related arrhythmias or heart failure **(Fig. 1D, E)**. We inspected three other *RBFOX2* alternative exons in unaffected fetal, adult, and DM1 heart tissues, however, their splicing pattern was unaltered during development, or in DM1 **(Supplementary Fig. 1F)**.

To determine if the increase in RBFOX2 protein abundance and change in its splicing were direct effects of the CTG expansion, we transfected HL-1 cardiac cell cultures with plasmids expressing zero (DT0) or 960 CTG repeats (DT960). Similar to DM1 heart tissues, transient transfection of HL-1 cells with the DT960 plasmid led to a significant elevation in RBFOX2 steady-state protein levels and a simultaneous shift in splicing from 43-nt to the 40-nt exon **(Fig. 1F-G and Supplementary Fig. 1G)**. To explore potential coupling between the increase in RBFOX2 protein and its isoform switch, we redirected *Rbfox2* splicing in HL-1 cells using splice-site (ss) blocking anti-sense oligonucleotides (ASO)^48^. Relative to control, HL-1 cells treated with ASO targeting the 5’ss of 43-nt exon caused a near complete switch to the 40-nt containing isoform but had little effect on RBFOX2 protein abundance **(Fig. 1H-I)**. These data indicate that increase in RBFOX2 and its splice isoform switch in DM1 are two separate events, and that upregulation of RBFOX2 protein is not a consequence of higher stability of the non-muscle isoform.

### Reduced expression of miRNAs de-represses RBFOX2 in DM1 cardiac cultures

Given that RBFOX2 protein levels decrease during postnatal heart development^23^ and the decrease is quickly reversed upon *Dicer* deletion in the adult myocardium^21^, we reasoned that RBFOX2 upregulation in DM1 might result from miRNA deregulation^20, 49, 50^. Consistent with this notion, we found conserved binding sites for a set of miRNAs—Let-7g/7b, miR-135a, miR-9, and miR-186—within the 3’-UTR of *Rbfox2* **(Fig. 2A)**, and expression of these miRNAs is reduced in DM1 heart tissue^20^. Transient expression of r(CUG)^960^ RNA was sufficient to downregulate all five miRNAs in HL-1 cells **(Fig. 2B)**; and likewise, transfections with Let-7g, miR-186, Let-7b, miR-9, and miR-135a mimics reduced RBFOX2 protein abundance compared to scrambled control or the non-targeting miR-30 mimic **(Fig. 2C)**, without affecting *Rbfox2* mRNA levels **(Supplementary Fig. 2A)**.

**Figure 2.**
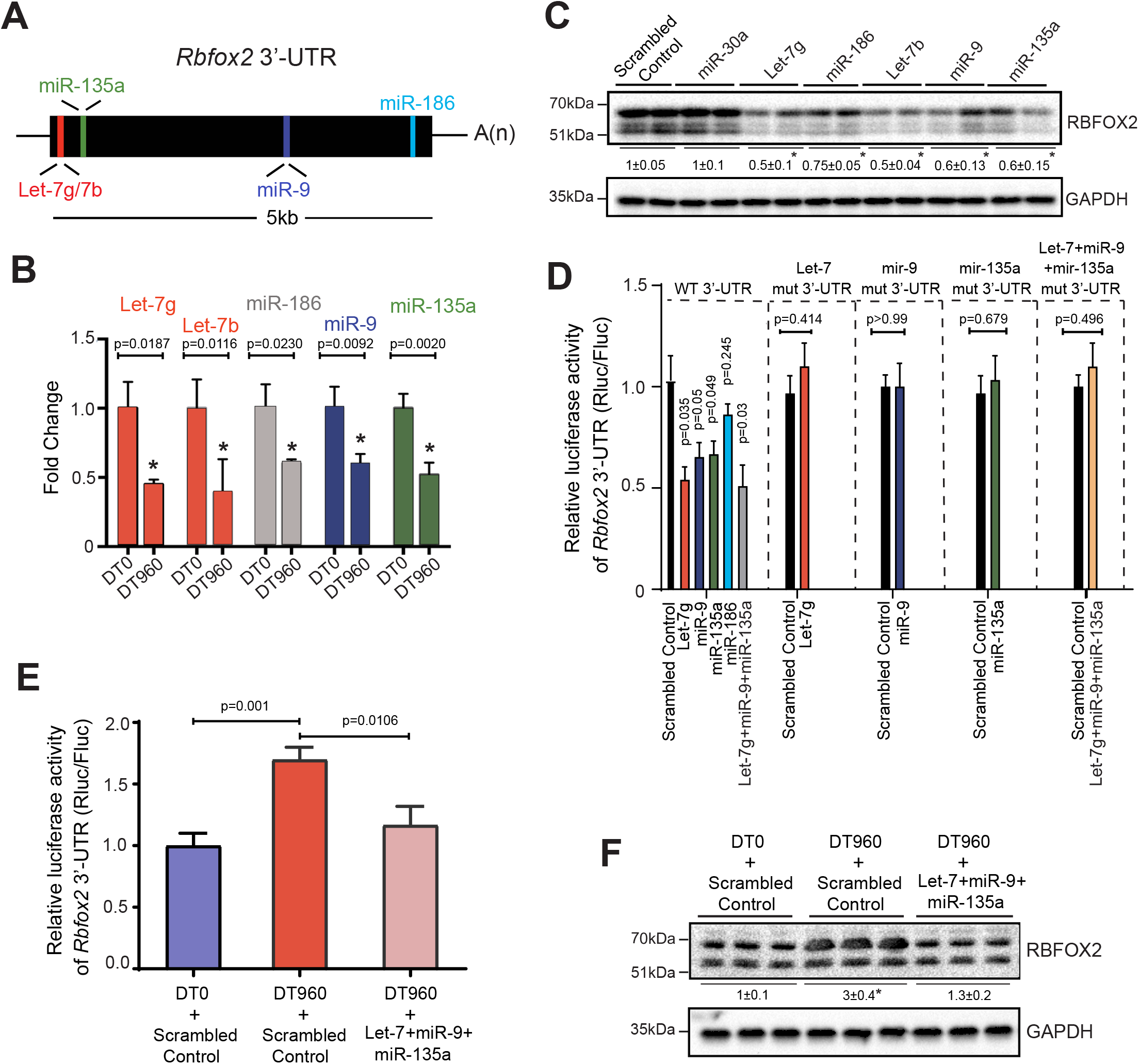
Reduced expression of miRNAs de-represses RBFOX2 in DM1 cardiac cultures. **(A)** Schematic of putative miRNAs targeting *Rbfox2* 3’-UTR that are downregulated in the hearts of DM1 patients ^20^. **(B)** Relative expression of indicated miRNAs in HL-1 cells following transfection with DT0 or DT960 plasmids for 48h. n=6 independent transfections. **(C)** Immunoblot analysis of RBFOX2 protein in HL-1 cells following treatment with scrambled control or indicated miRNA mimics. n=4 independent transfections. **(D)** Relative luciferase activity derived from *Rbfox2* 3’-UTR reporter transfected in HEK 293T cells following treatment with scrambled control or indicated miRNA mimics. Direct binding of miRNAs to the seed sequences in *Rbfox2* 3’-UTR was examined by co-transfecting indicated miRNA mimics and reporters with mutations that would disrupt the predicted miRNA interactions. n=4 independent transfections. **(E)** Relative luciferase activity derived from *Rbfox2* 3’-UTR reporter in HEK 293T cells, and **(F)** Immunoblot analysis of RBFOX2 protein in HL-1 cells after co-transfection with DT0 or DT960 plasmids, and scrambled control or a cocktail of indicated miRNA mimics. n=4 independent transfections. Data are mean ± s.d., and p-values were derived from a parametric t-test (two-sided, unpaired), with Welch’s correction.

To identify which of these miRNAs regulate RBFOX2 through direct interactions, we constructed dual luciferase reporters^51^ wherein the 3’-UTR of *Rbfox2* was cloned downstream from Renilla luciferase ORF, with or without mutations in seed sequences that would disrupt the predicted miRNA interactions **(Supplementary Fig. 2B)**. Co-transfection of HEK293T cells with the reporter plasmids and miRNA mimic(s) individually or in combination revealed that relative to control, Let-7g, miR-9, and miR-135a could each repress the luciferase (*Rluc/Fluc*) activity of wildtype, but not the mutant constructs **(Fig. 2D)**. Despite lowering RBFOX2 protein levels, miR-186, did not alter the reporter activity **(Fig. 2C, D)**. We next asked whether restoring the activity of select miRNAs can normalize RBFOX2 levels in a DM1 cardiac cell culture model. Owing to the reduction of *Rbfox2* targeting miRNAs, co-transfection of HL-1 cells with DT960, relative to DT0 plasmid, increased the *Rbfox2* 3’-UTR luciferase reporter activity **(Fig. 2B, E)**. Notably, supplementing the r(CUG)^960^ RNA expressing HL-1 cells with a cocktail of Let-7g, miR-9 and miR-135a mimics normalized not only the reporter activity **(Fig. 2E)** but also endogenous RBFOX2 protein levels **(Fig. 2F)**. *Rbfox2* mRNA levels were unchanged in these experiments **(Supplementary Fig. 2C)**. Altogether, these findings illustrate that multiple miRNAs directly or indirectly repress RBFOX2 protein expression in cardiac cells, their downregulation de-represses RBFOX2 in DM1 heart, and when supplemented, a mixture of miRNA mimics can reverse this de-repression in a DM1 cardiac cell culture model.

### CELF1 upregulation promotes the production of RBFOX2_40_ isoform in the heart

We next sought to address the mechanism(s) promoting generation of RBFOX2_40_ splice isoform in DM1. Because *DMPK* RNAs containing r(CUG)^exp^ repeats disrupt alternative splicing through MBNL sequestration and/or CELF1 upregulation, we assessed *Rbfox2* splicing pattern in the hearts of *Mbnl1*^ΔE3/ΔE3^, tetracycline (*tet*)-inducible TRE-CELF1; MHCrtTA, and littermate control mice^23, 27^. Although CELF1 overexpression in the adult heart did not perturb RBFOX2 protein abundance **(Supplementary Fig. 3A)**, it evoked a significant shift from 43-nt to 40-nt exon within *Rbfox2* transcripts, mimicking the splicing pattern observed in DM1 heart tissue **(Fig. 3A, B)**. Deletion of MBNL1 had no effect on *Rbfox2* splicing. We analyzed published time-series RNA-seq data following CELF1 overexpression in mouse hearts^25^ and found a progressive shift in *Rbfox2* splicing, which was detected as early as 24h after CELF1 induction **(Fig. 3C)**. Furthermore, forced expression of CELF1 in HL-1 cells was sufficient to induce a switch from 43-nt to 40-nt exon without affecting RBFOX2 protein levels **(Fig. 3D, E and Supplementary Fig. 3B)**. As expected, r(CUG)^960^ RNA expression resulted in CELF1 upregulation, and a consequent shift in *Rbfox2* splicing in these cells **(Fig. 3F, G)**. To determine whether increased CELF1 levels were required to generate the RBFOX2_40_ isoform in DM1, we used siRNA(s) to offset CELF1 upregulation in r(CUG)^960^ RNA expressing HL-1 cells. Relative to scrambled control, *Celf1*-targeting siRNAs normalized the CELF1 protein levels and partially abrogated the r(CUG)^960^ RNA-induced switch in *Rbfox2* splicing **(Fig. 3F, G)**. These data imply that CELF1 promotes the production of non-muscle RBFOX2_40_ isoform in the heart, and that high CELF1 activity is in part required for producing this isoform in a DM1 cardiac cell culture model.

**Figure 3.**
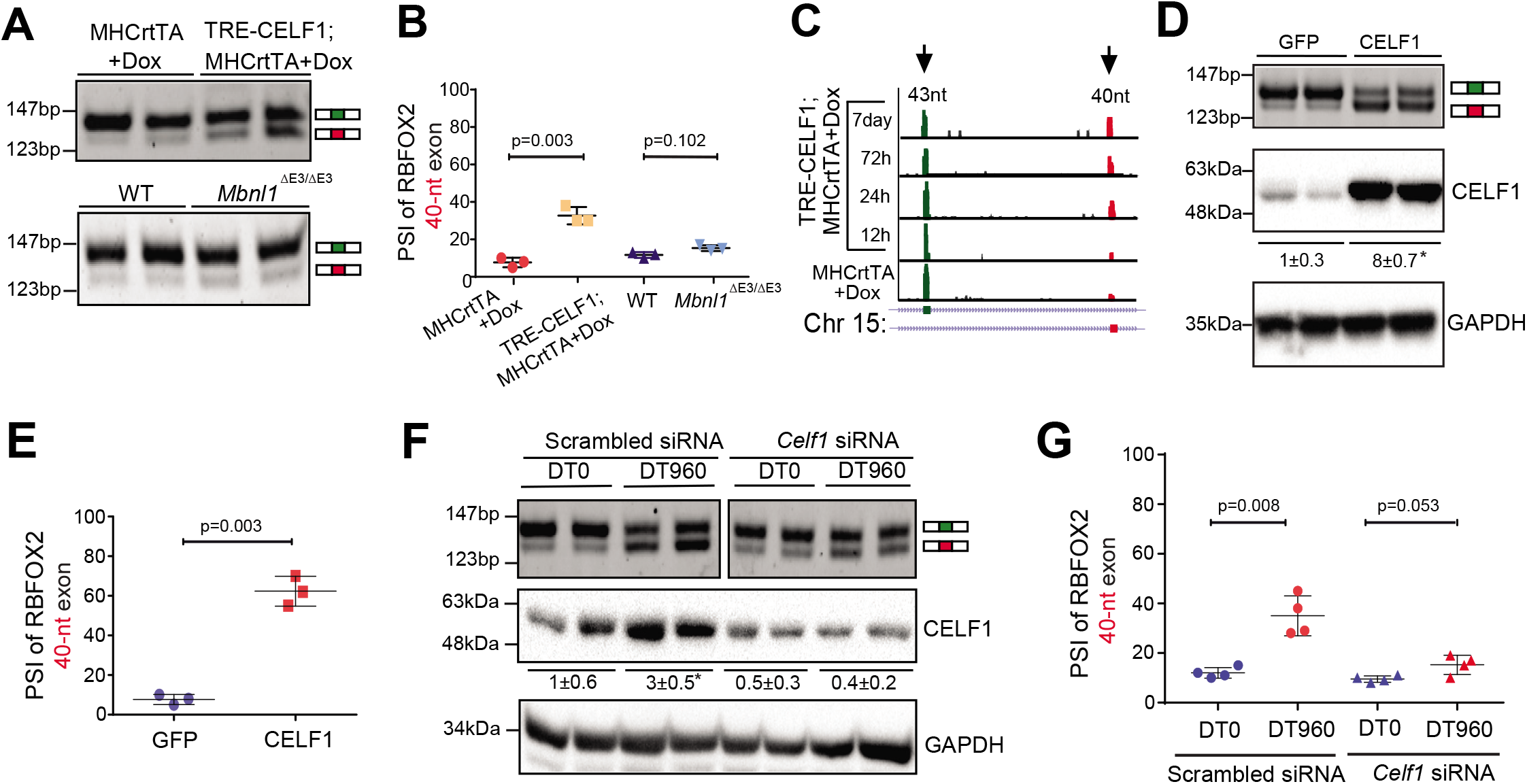
CELF1 upregulation promotes the production of non-muscle RBFOX2_40_ isoform in the heart. **(A)** RT-PCR analysis of *Rbfox2* 43-nt and 40-nt exons in hearts of *tet*-inducible, heart-specific CELF1 bitransgenics (TRE-CELF1; MHCrtTA) and littermate control mice induced with 6g/kg Dox for ten days, as well as in hearts of wildtype (WT) and *Mbnl1*^ΔE3/ΔE3^ mice. **(B)** PSI values of *Rbfox2* 40-nt exon from the indicated genotypes in **A**. n=4 mice for each genotype. **(C)** RNA-seq read density across *Rbfox2* 43-nt and 40-nt exons from mouse hearts following CELF1 overexpression at indicated time points ^25^. **(D)** RT-PCR analysis of *Rbfox2* 43-nt and 40-nt exons (top), and immunoblot analysis of CELF1 protein normalized to GAPDH (bottom) in HL-1 cells following infection with GFP or CELF1 expressing adenoviruses. **(E)** PSI values of *Rbfox2* 40-nt exon from **D**. n=4 independent infections. **(F)** RT-PCR analysis of *Rbfox2* 43-nt and 40-nt exons (top) and immunoblot analysis of CELF1 protein normalized to GAPDH (bottom) in HL-1 cells after co-transfection with scrambled control or *Celf1* targeting siRNA(s), and DT0 or DT960 plasmids. **(G)** PSI values of *Rbfox2* 40-nt exon from **F**. n=4 independent transfections. All data are mean ± s.d., and p-values were derived from a parametric t-test (two-sided, unpaired), with Welch’s correction.

### RBFOX2_40_ isoform induces DM1-like cardiac pathology in mice

Next, we characterized the functional consequences of RBFOX2_40_ isoform in DM1 cardiac pathogenesis. To distinguish the effects of overexpression from splice isoform switch, we generated two separate mice models: a *tet*-inducible transgenic mouse model that overexpresses FLAG-tagged RBFOX2_40_ isoform in the adult heart; and a CRISPR/Cas9 derived *Rbfox2* 43-nt exon deletion mouse model **(Supplementary Fig. 4A-D)**. When induced with doxycycline (Dox), the hemizygous TRE-RBFOX2_40_; MHCrtTA bitransgenic mice produced approximately 8-fold higher RBFOX2 protein compared to the uninduced littermate controls **(Fig. 4A, B and Supplementary Fig. 4E)**. On the other hand, homozygous deletion of 43-nt exon by CRISPR/Cas9 in *Rbfox2*^Δ43/Δ43^ mice redirected splicing to 40-nt exon generating the non-muscle isoform in cardiac and skeletal muscles, but without any change in RBFOX2 mRNA or protein abundance **(Fig. 4C, D and Supplementary Fig. 4E, F)**.

**Figure 4.**
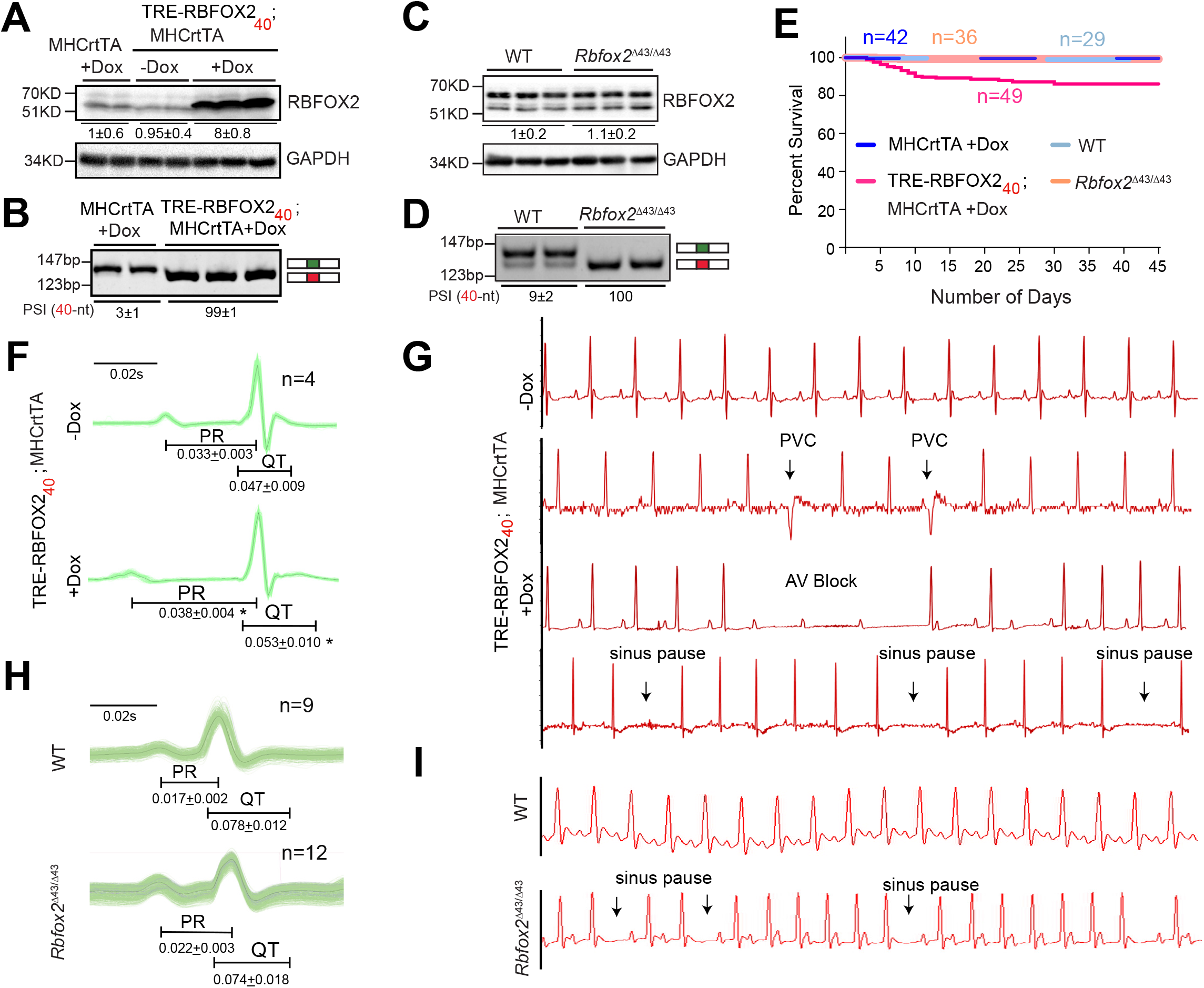
RBFOX2_40_ isoform expression induces DM1-like cardiac pathology in mice. **(A)** Immunoblot analysis of RBFOX2 and GAPDH proteins, and **(B)** RT-PCR analysis of *Rbfox2* 43-nt and 40-nt exons in the hearts of hemizygous MHCrtTA and TRE-RBFOX2_40_; MHCrtTA bitransgenic mice fed regular or 0.5g/kg Doxycycline (Dox) containing Chow for 3 days. Quantifications of relative band intensities of RBFOX2 normalized to GAPDH, and the PSI values of *Rbfox2* 40-nt exon are shown below the gel images. n=5 mice for each genotype. **(C)** Immunoblot analysis of RBFOX2 and GAPDH proteins, and **(D)** RT-PCR analysis of *Rbfox2* 43-nt and 40-nt exons in the hearts from wildtype (WT) and *Rbfox2*^Δ43/Δ43^ mice. Quantifications of relative band intensities of RBFOX2 normalized to GAPDH, and the PSI values of *Rbfox2* 40-nt exon are shown below the gel images. n=4-6 mice for each genotype. **(E)** Kaplan-Meier survival curves of mice from indicated genotypes fed regular or 0.5g/kg Dox containing Chow for 45 days. **(F)** ECG analysis of ambulatory TRE-RBFOX2_40_; MHCrtTA bitransgenic mice 48h before and 9 days after RBFOX2_40_ overexpression with 0.5g/kg Dox containing Chow. **(G)** Representative ECG traces from F recorded before and after RBFOX2_40_ overexpression. PVC: premature ventricular contraction, AV block: atrioventricular block. **(H)** Surface ECG analysis of WT and *Rbfox2*^Δ43/Δ43^ mice. **(I)** Representative ECG traces from H. PR, and QT intervals in **F** and **H** are indicated below the traces. All data are mean ± s.d., and p-values were derived from a parametric t-test (two-sided, paired for **F** and unpaired for **H**), with Welch’s correction.

Strikingly, ~20% of TRE-RBFOX2_40_; MHCrtTA bitransgenic mice died within two to three weeks of transgene expression whereas no mortality was seen in *Rbfox2*^Δ43/Δ43^ or littermate control mice **(Fig. 4E)**. Because no apparent differences in heart-to-body weight ratios, atrial and ventricular size or histological abnormalities such as myofiber disarray, inflammation, and fibrosis were evident **(Supplementary Fig. 4G-I)**, we suspected that sudden death following RBFOX2_40_ overexpression in the heart might be related to electrical phenotypes. Hence, we performed ECG telemetry on un-anaesthetized TRE-RBFOX2_40_; MHCrtTA bitransgenic mice—continuously recording their ECG waveforms—before (for 48h) and after (for nine days) of Dox administration. In addition, we also did surface ECGs on a separate cohort of wildtype, *Rbfox2*^Δ43/Δ43^, TRE-RBFOX2_40_; MHCrtTA bitransgenics and littermate control mice following three days of Dox administration.

Compared to the baseline recordings, all mice following overexpression of RBFOX2_40_ isoform exhibited a significant increase in PR interval, indicating slowed conduction from the atria to the ventricles and prolonged QT intervals, indicating prolonged ventricular repolarization **(Fig. 4F and Supplementary Fig. 5A, B)**. Importantly, while the uninduced mice displayed a normal sinus rhythm, progressive increases in the number of pauses and arrhythmic events were detected following RBFOX2_40_ overexpression. Heart rate variability and sinus pauses indicated sinus node dysfunction, sometimes resulting in ventricular escape beats. Additionally, we observed premature ventricular contractions and frequent atrioventricular conduction blocks—evidenced by non-conducting P-waves **(Fig. 4G)**. Furthermore, nine days after transgene induction, RBFOX2_40_ overexpressing mice showed a high variability in RR interval duration **(Supplementary Fig. 5C)**. Echocardiography studies revealed that abnormal conductivity in these mice is accompanied by reduced ejection fraction, fractional shortening and systolic/diastolic alterations **(Supplementary Fig. 5D)**. Thus, overexpression of RBFOX2_40_ isoform in mouse heart recapitulates DM1 cardiac phenotypes including conduction delay, spontaneous episodes of arrhythmias and contractile dysfunctions.

Although the electrophysiological defects in *Rbfox2*^Δ43/Δ43^ mice were milder relative to RBFOX2_40_ overexpressing mice, ECG analysis of *Rbfox2*^Δ43/Δ43^ mice reproduced certain pathologic features of DM1. For instance, we noted a clear trend of prolonged PR interval with narrower QRS morphology in *Rbfox2*^Δ43/Δ43^ mice **(Fig. 4H)**, which reflects supraventricular origin of QRS complex, and suggests that arrhythmias in these mice originate above or within the bundle of His. The slow conductivity was also accompanied by an increase in sinus pauses, atrioventricular blocks and ventricular electric events **(Fig. 4I)**. Histological and echocardiography studies, however, showed no differences in the systolic/diastolic ventricular dimensions or contractility including fractional shortening between *Rbfox2*^Δ43/Δ43^ and wildtype mice **(Supplementary Fig. 4I, 5D)**. Collectively, these results demonstrate that inappropriate expression of the non-muscle RBFOX2_40_ isoform in mouse heart disrupts the cardiac conduction, rhythm and function; and analogous to DM1 patients, triggers progressive heart blocks and sudden death in a subset of mice.

### RBFOX2_40_ isoform driven transcriptome alterations in DM1 heart tissue

To further investigate the mechanism(s) by which RBFOX2_40_ isoform provokes DM1-like cardiac pathology, we deep-sequenced poly(A) selected RNAs prepared freshly from the ventricular cardiomyocytes isolated from Dox-induced TRE-RBFOX2_40_; MHCrtTA and littermate control mice **(Supplementary Table 2 and Supplementary Fig. 6A, B)**. In order to minimize any secondary effects of arrhythmias on the transcriptome, that are not directly dependent on RBFOX2_40_, we induced the mice for only 72h. Remarkably, even a short-term induction of RBFOX2_40_ transgene produced extensive changes in mRNA splicing [3,144 events within 1,734 mRNAs with ΔPSI>20%; false discovery rate (FDR)<0.10] and abundance [2,679 mRNAs with fold change>2.0; FDR<0.05] **(Fig. 5A and Supplementary Fig. 6C)**. The differentially expressed mRNAs following RBFOX2_40_ overexpression were enriched (*P_adj_* <0.05) in gene ontologies that grouped into major functional clusters especially cardiac conduction, immune response, cell proliferation, apoptosis, and metabolic processes, whereas the gene set with altered splicing formed clusters related to transcriptional regulation, mRNA processing, cytoskeletal organization and transport functions **(Supplementary Fig. 6D)**.

**Figure 5.**
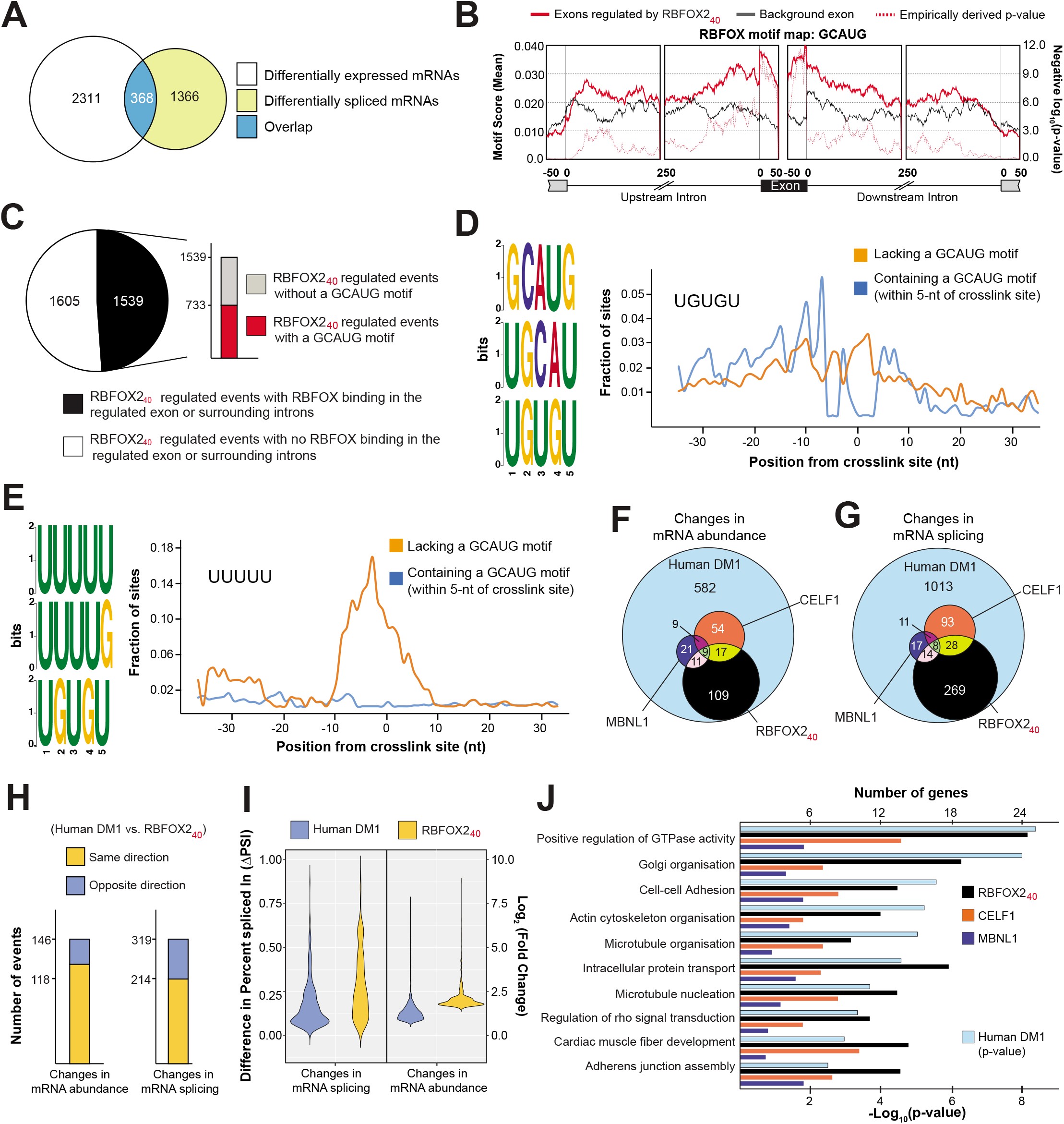
RBFOX2_40_ isoform driven transcriptome alterations in DM1 heart tissue. **(A)** Overlap of differentially expressed (p<0.05, Wald test as described DESeq2; Log2[Fold Change]>1, and TPM>4) with differentially spliced mRNAs (p<0.05, FDR<0.10 adjusted for multiple testing, Difference in Percent Spliced Index [ΔPSI]≥20%, and Junction Counts≥10) in cardiomyocytes isolated from hemizygous MHCrtTA, and TRE-RBFOX2_40_; MHCrtTA bitransgenic mice after administration of 0.5g/kg Dox containing Chow for 3 days. **(B)** Position and relative enrichment of RBFOX-binding motif near RBFOX2_40_ regulated cassette exons. **(C)** Breakup of 3144 RBFOX2_40_ regulated splicing events in cardiomyocytes with or without RBFOX2_40_ binding peaks and a GCAUG motif within those peaks from iCLIP data in mouse brain ^46^. **(D-E)** Top 3 pentamers enriched within the binding peaks near RBFOX2_40_ regulated exons with **(D)** or without **(E)** GCAUG motifs. Fractional enrichments of **(D)** UGUGU and **(E)** UUUUU sequences aligning at each nucleotide relative to the RBFOX2_40_ crosslink site are also plotted. Overlap of **(G)** mRNA abundance, and **(F)** alternative splicing changes among cardiac transcriptomes of DM1 patients ^52^, RBFOX2_40_ overexpressing, CELF1 overexpressing ^25^, and *Mbnl1*^ΔE3/ΔE3^ mice ^25^. **(H)** Directionality of mRNA abundance and alternative splicing changes in DM1 patient hearts and RBFOX2_40_ overexpressing cardiomyocytes. **(I)** Distribution of changes in mRNA abundance and splicing events co-regulated in DM1 patient hearts and RBFOX2_40_ overexpressing cardiomyocytes. **(J)** p-values of top Gene Ontology terms for alternatively spliced mRNAs in DM1 patient hearts (hypergeometric test; Benjamini method for multiple testing), and corresponding numbers of genes for each category that are similarly misspliced in the cardiomyocytes of RBFOX2_40_ overexpressing, and hearts of CELF1 overexpressing, or *Mbnl1*^ΔE3/ΔE3^ mice.

Consistent with earlier studies^39, 40^, the RBFOX2 binding motif GCAUG was significantly overrepresented in the exonic and surrounding intronic regions of RBFOX2_40_-regulated exons **(Fig. 5B)**. We next integrated the RBFOX2_40_ overexpression RNA-seq data with previously published RBFOX2_40_ iCLIP data from mouse brain^46^. Overlay of these two datasets revealed that ~50% of misspliced pre-mRNAs had a robust RBFOX2-binding cluster in the regulated exon or surrounding introns, but only half of those clusters contained a GCAUG motif **(Fig. 5C)**. Because the RBFOX2_40_ isoform in LASR complex can be recruited to RNA directly or indirectly through binding of hnRNP proteins^47^, we searched for pentamers near RBFOX2_40_-iCLIP crosslink sites bearing a GCAUG motif, or not **(Supplementary Fig. 6E, F)**. Besides the expected overrepresentation of GCAUG sequences, a clear enrichment of potential hnRNP M-binding motif UGUGU was noted directly next to the GCAUG containing crosslink sites **(Fig. 5D and Supplementary Fig. 6E)**. On the contrary, the polyU-rich hnRNP C-binding motifs were the most enriched pentamers detected adjacent to non-GCAUG containing crosslink sites **(Fig. 5E and Supplementary Fig. 6F)**. Thus, our cardiomyocyte data are in agreement with previous studies^46, 47^, that in addition to direct interactions, the non-muscle RBFOX2_40_ isoform can target pre-mRNAs through RNA contacts of hnRNP M and C components of the LASR complex.

To determine the respective contribution of RBFOX2_40_ overexpression towards DM1—in relation to misregulation of MBNL and CELF proteins—we analyzed published cardiac transcriptomes of DM1 patients^52^, CELF1 overexpressing and *Mbnl1*^ΔE3IΔE3^ mice^25^, and compared them with the cardiac transcriptome of RBFOX2_40_ overexpressing mice. Amongst the three mice models, RBFOX2_40_ overexpression reproduced the highest mRNA abundance and splicing defects observed in heart tissue of DM1 patients **(Fig. 5F, G)**. A large proportion of these mRNA abundance (74%) and splicing (67%) defects were shared in directionality **(Fig. 5H)**; however, relative to DM1 hearts, the overall magnitude of change was higher in RBFOX2_40_ overexpressing mouse cardiomyocytes **(Fig. 5I)**. For the coregulated sets of splicing events, we found a positive correlation of CELF1 with RBFOX2_40_ and a negative correlation of MBNL1 with CELF1 and RBFOX2_40_ proteins **(Supplementary Fig. 6G)**. Thus, our analysis confirmed the antagonistic splicing regulation by CELF1 and MBNL1 proteins^23, 25^ while revealing a synergistic mode of regulation between RBFOX2_40_ and CELF1 proteins. The involvement of RBFOX2_40_ in DM1 cardiac pathology was further strengthened by the large overlap of gene ontologies between DM1 patients and RBFOX2_40_ overexpressing mice **(Fig. 5J)**. Altogether, our comparative transcriptomic findings illustrate that upregulation of non-muscle RBFOX2_40_ isoform in DM1 alters the mRNA abundance and splicing of a large group of genes via direct and/or indirect pre-mRNA interactions through the LASR complex.

### Structure-function analysis of DM1-associated alternatively spliced isoforms of voltage-gated ion channels

Further analysis of the RBFOX2_40_-driven splicing alterations in DM1 identified twenty-two genes encoding components of conduction system and/or contractile apparatus—many of which are related to the NCBI MeSH terms: “Ventricular Tachycardia,” “Brugada and Long QT Syndromes,” “Atrial Fibrillation” and “Sudden Cardiac Death” **(Fig. 6A)**. Of these, we elected to focus on *SCN5A, CACNA1C*, and *KCND3* because they encode the major voltage-gated sodium, calcium, and potassium channels in the heart respectively; and they switch to producing their pro-arrhythmic splice isoforms^52–55^ in DM1 patients. Additionally, multiple genome-wide association studies have identified variants of *SCN5A* and *KCND3* among genetic determinants of prolonged PR interval and QRS duration^56, 57^, which are distinctive features of cardiac-conduction delay in DM1 patients.

**Figure 6.**
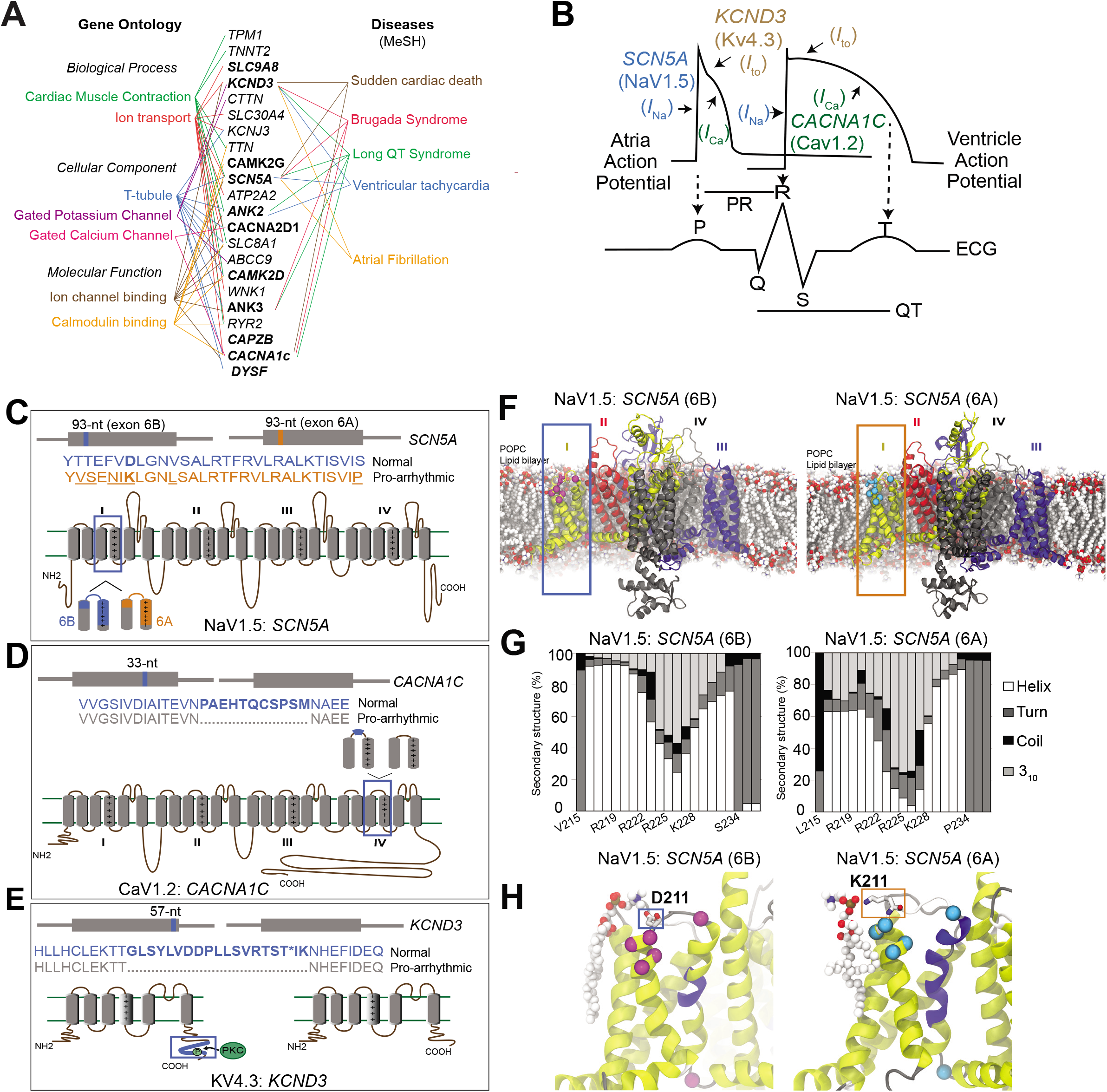
Structure-function analysis of normal and DM1-related splice isoforms of voltage-gated sodium, calcium, and potassium channels. **(A)** The subset of cardiac arrhythmia-associated genes misspliced following RBFOX2_40_ overexpression in cardiomyocytes with their respective Gene Ontology terms (Left) and MeSH disease headings (Right). Genes misspliced similarly in the hearts of DM1 patients are highlighted in bold letters. **(B)** Schematic representation of action potentials from atria and ventricles, and their correlation with the surface electrocardiogram (ECG). The sodium channel *SCN5A* (Na_V_1.5) mediated *I_na_* current, calcium channel *CACNA1C* (Ca_V_1.2) mediated *I_ca_* current, and potassium channel *KCND3* (K_V_4.3) mediated *I_to_* current with their respective functions in propagation and duration of a typical action potential are depicted. Cartoons representing topology, relative locations and amino acid sequences for the two alternatively spliced exons in **(C)** *SCN5A*, **(D)** *CACNA1C* and **(E)** *KCND3* proteins. **(F)** Molecular dynamics (MD) simulation showing modelled structure of Na_V_1.5:SCN5A (6B) and (6A) exon containing isoforms embedded in a 1-palmitoyl-2-oleoyl-sn-glycero-3-phosphocholine (POPC) lipid bilayer. Domains I-IV of the channel are represented by yellow, red, black and blue colors respectively. Differences in protein sequence are highlighted on the structure by magenta/cyan spheres and sticks. **(G)** Secondary structure content of domain I segment S3-S4 observed in MD simulations for the two isoforms. **(H)** Representative snapshots of MD simulations highlighting differential interactions of D211 in 6B (Left) and K211 in 6A (Right) with the lipid head group. The non-conserved residues between the two isoforms are shown in Van der Waals (VdW) representation. The protein is colored based on the secondary structure, yellow: α-helix, blue: 3_10_-helix, gray: turn, white: coil.

For each cardiac cycle, when the pacemaker cells in the sinoatrial node fire, the action potential spreads through the atrial and ventricular myocardium via gap junctions that allow it to propagate along the myocytes. Once the action potential arrives, activation the membrane potential of myocytes increases, which activates the voltage-gated SCN5A sodium channels (Na_V_1.5) to mediate a rapid influx of Na^+^ ions (*I*_Na_) across the cell membrane, resulting in a fast depolarization phase^58^ **(Fig. 6B)**. As the membrane potential continues to increase, the voltage-gated L-type CACNA1C calcium channels (Ca_V_1.2) open, and Ca^2+^ ions flow into the cells as an inward current (*I*_Ca_), causing additional release of Ca^2+^ from the sarcoplasmic reticulum, which is crucial for coupling the electrical excitation to the physical contraction of the myocyte^59^ **(Fig. 6B)**. After the membrane potential peaks, the voltage-gated KCND3 potassium channels (K_V_4.3) open and potassium ions flow out of the cells generating a rapidly inactivating transient outward current (*I*_to_), which is important for the eventual repolarization of cardiac myocytes^60^ **(Fig. 6B)**. Thus, proper impulse propagation and duration of the action potential in heart is highly contingent on the precise function of these ion channels.

Structurally, the voltage-gated ion channels are made up of either four separate subunits (K^+^ channels) or one polypeptide with four homologous domains (Na+ and Ca^2+^ channels) **(Fig. 6C-E)**. Each subunit or domain has six transmembrane segments (S1-S6) and a pore loop. The voltage sensing transmembrane element in SCN5A lies within segments S3 and S4, and that of domain I is encoded by a pair of conserved, developmentally regulated, mutually exclusive exons that are each 93-nts long. The fetal exon 6A differs from the adult exon 6B by 31-nts creating seven amino acid substitutions in Nav1.5 protein^61^, which includes replacement of a positively charged lysine (K211) in exon 6A with a negatively charged aspartate (D211) in exon 6B **(Fig. 6C)**. Because a shift from exon 6B to 6A is linked to DM1-associated cardiac-conduction delay and heart arrhythmias^52, 54^, we investigated functionally relevant structural and dynamical differences between the two alternatively spliced Nav1.5 isoforms using molecular dynamics (MD) simulations. Given that the three-dimensional structure of Nav1.5 channel is unavailable, we built homology models of the exon 6A or exon 6B containing human Na_v_1.5 isoforms using existing structures of the activated sodium channels Na_v_1.4 and Na_v_Pas^62, 63^. These models were then used in subsequent MD simulations, where three independent 100ns-long equilibrium simulations were carried out on 6A or 6B containing Na_v_1.5 isoforms. In all simulations, the channels were embedded in a lipid bilayer composed of 1-palmitoyl-2-oleoyl-sn-glycero-3-phosphocholine (POPC) **(Fig. 6F)**.

Our simulations revealed that in both channels, the S4 helix of domain I largely maintains its α-helical secondary structure **(Fig. 6G)**, but with different tendencies to form a more tightly wound, longer, and thinner 3_10_-helix at different segments of S4. Analysis of structural dynamics during the simulations showed that, while similar propensities for 3_10_-helix formation were evident in the middle of S4 helix for both 6A and 6B isoforms, the former exhibited a higher 3_10_-helix content near the top portion of S4, which can facilitate the downward S4 transition from the activated (up) state^64^ **(Fig. 6G, H)**. Frequent transitions observed between α- and 3_10_-helix further indicate that the activated state of S4 in the 6A isoform is destabilized and it may deactivate more readily due to a reduction in energy barrier, which is consistent with a depolarized shift in the voltage-dependence of channel activation for the pro-arrhythmic 6A isoform^52, 54^. Additionally, we note that although D211 in 6B and K211 in 6A are both positioned close to the lipid headgroups, unlike D211, which preferably interacts with the choline moiety at a slightly higher position with respect to the lipid bilayer, K211—due to its longer side-chain—can reach down into the headgroup region and coordinate closely with the nearby phosphate group of POPC lipids **(Fig. 6H, Supplementary Fig. 7)**. The K211D variant shifts the voltage-dependence of activation of the 6A containing Nav1.5 isoform back towards the hyperpolarized direction^65^. We propose that the observed persistent lipid-interaction of K211 in 6A isoform is crucial for stabilizing a 3_10_-helical conformation near the top portion of S4 to allow rapid deactivation that would slow the activation kinetics **(Supplementary video 1, 2)**. Indeed, forced expression of the exon 6A containing Na_V_1.5 isoform in adult mouse heart reduces excitability, slows conduction velocity, prolongs PR and QRS intervals, and increases susceptibility to arrhythmias^52, 54^.

The primary cardiac isoform of the α1C subunit of L-type calcium channel *CACNA1C* (Ca_V_1.2) includes a 33-nt alternative exon that increases the length of the linker separating the S3 and S4 segments of domain IV^66^ **(Fig. 6D)**. Inclusion of 11 amino acids encoded by this exon changes the activation potential of Ca_V_1.2 channel, requiring larger depolarization of membrane potential to open^67^. Of note, targeted deletion of the 33-nt exon from *Cacna1c* gene in mice was recently shown to prolong the ventricular cardiomyocyte action potential duration and cause premature ventricular contractions, tachycardia, and increased QT interval^53^. Likewise, the cardiac isoform of the a subunit of potassium channel *KCND3* (K_V_4.3) includes a 57-nt alternative exon, which encodes 19 amino acids at the C-terminal end **(Fig. 6E)**. Amongst these amino acids is a conserved threonine residue, which is phosphorylated by protein kinase C to facilitate regulation of transient outward current *I*_to_^68^. The K_V_4.3 isoform lacking the 57-nt exon has been shown to promote faster inactivation of the channel, which slows down the ion flux and lengthens the duration of action potential—particularly affecting the early recovery and plateau phases of repolarization^69^.

### RBFOX2_40_ expression in the heart reproduces DM1-related missplicing of voltage-gated sodium and potassium channels

Owing to such drastic effects of ion channel variants on cardiac rhythm and conductivity, we assessed their splicing patterns in heart tissues of unaffected and DM1 individuals as well as in wildtype, *Rbfox2*^Δ43/Δ43^, TRE-RBFOX2_40_; MHCrtTA bitransgenics and littermate control mice. Notably, the indicated exons in *SCN5A* and *KCND3* ion channel transcripts were similarly misspliced in DM1 patients and RBFOX2_40_ overexpressing mice **(Fig. 7A)** but not in individuals diagnosed with other forms of arrhythmias **(Supplementary Fig. 8A)**. We noticed that *Scn5a* and *Kcnd3* were also misspliced in the hearts of *Rbfox2*^Δ43/Δ43^ mice, albeit to a lesser extent **(Fig. 7A)**, which might explain their milder arrhythmia phenotypes compared to RBFOX2_40_ overexpressing mice. Whereas the 33-nt exon in *CACNA1C* transcripts was predominantly skipped in DM1 patients, it was not skipped in the RBFOX2_40_ overexpressing or *Rbfox2*^Δ43/Δ43^ mice **(Fig. 7A)** likely because RBFOX proteins stimulate the inclusion of this exon^47^. Additionally, we detected large clusters of RBFOX2_40_ footprints upstream of the *Kcnd3* alternative exon in the iCLIP data^46^, indicating that the non-muscle RBFOX2_40_ isoform directly binds these intronic regions and, when expressed at high levels, causes skipping of the downstream exon **(Fig. 7B)**. As expected, no RBFOX2_40_ binding on the *Scn5a* transcript was detected in the brain iCLIP dataset because *Scn5a* is a cardiac-specific sodium channel and is not expressed in the brain **(Fig. 7B and Supplementary Fig. 8B, C)**.

**Figure 7.**
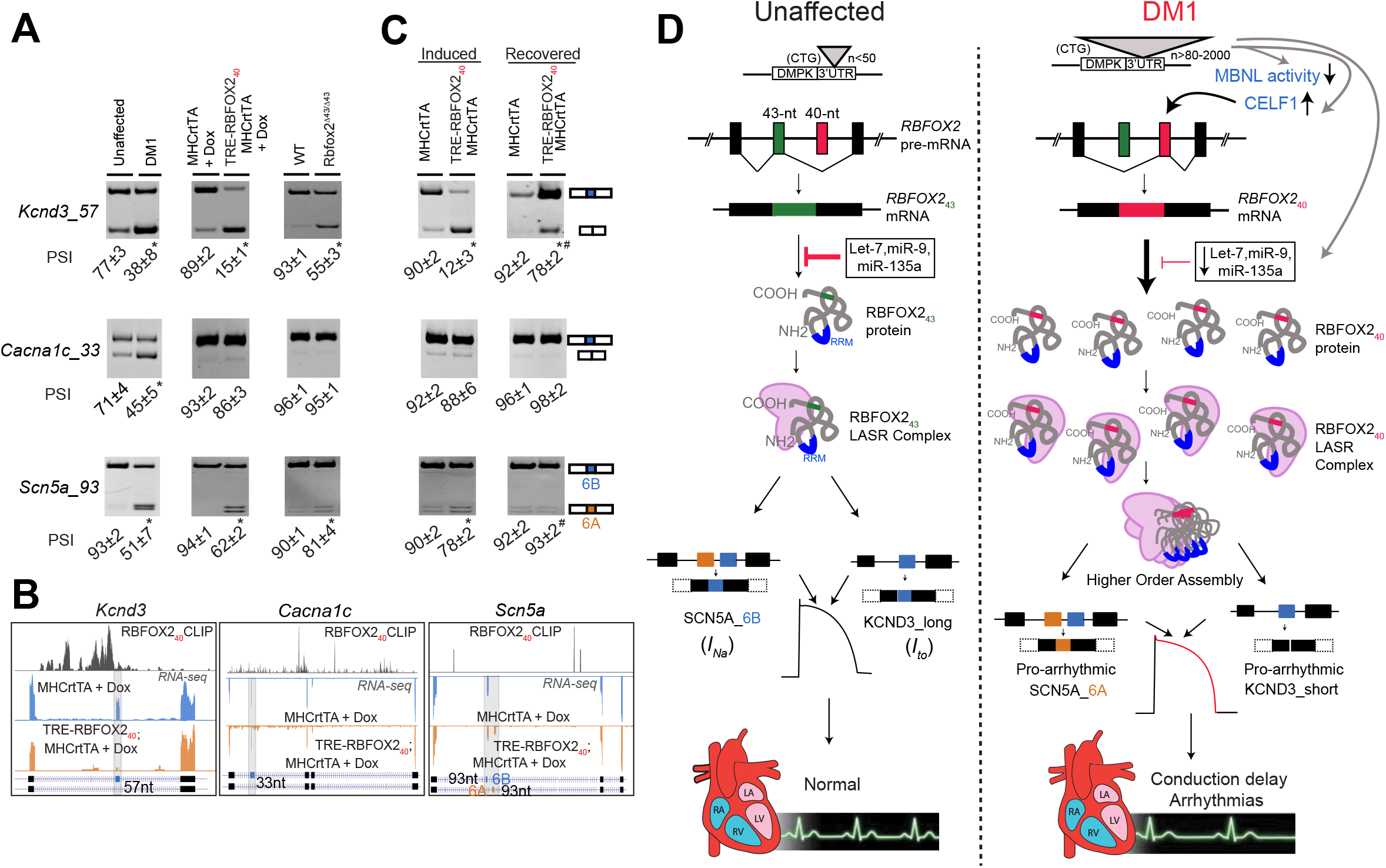
RBFOX2_40_ isoform expression in the heart reproduces DM1-related missplicing of voltage-gated sodium and potassium channels. **(A)** RT-PCR analysis comparing the inclusion of 57-nt exon in *KCND3*, 33-nt exon in *CACNA1C*, and 93-nt mutually exclusive exons (6A and 6B) in *SCN5A* transcripts in the hearts of unaffected individuals, DM1 patients, hemizygous MHCrtTA, and TRE-RBFOX2_40_; MHCrtTA bitransgenics induced with 0.5g/kg Dox for 3 days, as well as wildtype (WT) and *Rbfox2*^Δ43/Δ43^ mice. n=3 for all human and mouse samples. (B) UCSC genome browser snapshots demonstrating RBFOX2_40_ footprints and splicing patterns of indicated alternative exons in *Kcnd3, Cacna1c*, and *Scn5a* transcripts from 0.5g/kg Dox-induced TRE-RBFOX2_40_; MHCrtTA bitransgenics (Orange panel) and littermate control mice (blue panel). X-axis: the position of RBFOX2_40_ iCLIP data from mouse brain ^46^ (top track), or RNA-seq reads across indicated transcripts (bottom two tracks). Y-axis: normalized read density scaled for each track. **(C)** RT-PCR analysis of *Kcnd3, Cacna1c*, and *Scn5a* alternative exons in the hearts of TRE-RBFOX2_40_; MHCrtTA and littermate control mice induced with 0.5g/kg Dox for 24h followed by a 3-day recovery period. n=3 mice for each genotype/condition. All data are mean ± s.d., and p-values were derived from a parametric t-test (two-sided, unpaired), with Welch’s correction for **A**, and from one-way ANOVA plus Dunnett’s post-hoc test for **C**. *p<0.05 (between MHCrtTA and TRE-RBFOX2_40_; MHCrtTA), and #p<0.05 (between TRE-RBFOX2_40_; MHCrtTA (induced) and TRE-RBFOX2_40_; MHCrtTA (recovered). **(D)** Working model for the generation and function of non-muscle RBFOX2_40_ isoform in DM1 cardiac arrhythmogenesis.

To further investigate whether missplicing of ion channels following RBFOX2_40_ overexpression can be reversed, we first induced TRE-RBFOX2_40_; MHCrtTA mice with Dox for 24h and then silenced the transgene in a subset of mice by withdrawing Dox for a period of 3 days. In the induced mice, expression of RBFOX2_40_ non-muscle isoform was rapidly increased **(Supplementary Fig. 8D, E)** leading to a switch from exon 6B to 6A in *Scn5a* and skipping of the 57-nt exon in *Kcnd3* transcripts **(Fig. 7C)**. Turning off the transgene led to a marked reduction in RBFOX2_40_ levels **(Supplementary Fig. 8D, E)**, and near-complete restoration of both *Scn5a* and *Kcnd3* splicing to their normal patterns **(Fig. 7C)**. Overall, these data demonstrate that the non-muscle RBFOX2_40_ isoform directs the production of pro-arrhythmic splice variants of sodium and potassium channels that elicit altered activation-deactivation kinetics, rates of ion transport, and electrophysiological properties. Furthermore, quick reversal of ion channel missplicing upon silencing of RBFOX2_40_ transgene suggests that eliminating the aberrant RBFOX2_40_ isoform from heart might mitigate the DM1 cardiac phenotypes.

## DISCUSSION

Cardiac arrhythmias are a prominent cause of mortality in DM1 patients^9–11^; but, our understanding of the molecular events triggering electrophysiological abnormalities in DM1 heart is limited. In this study, we provide multiple lines of evidence that inappropriate expression of non-muscle RBFOX2_40_ isoform in DM1 heart leads to cardiac conduction delay, atrioventricular heart blocks, and spontaneous arrhythmogenesis. RBFOX family represents a multifunctional group of sequence-specific RNA binding proteins that are critical regulators of splicing in multiple tissues including skeletal muscle, heart, and brain^23, 36, 37, 40, 44, 70–73^. Loss-of-function mutations in *RBFOX2* gene have been associated with congenital heart disease in humans^41–43^; whereas RBFOX2 activity in diabetic hearts is inhibited due to upregulation of its dominant negative splice isoform^45, 74^. Recently, RBFOX1 and RBFOX2 proteins were shown to compete with MBNL1 for binding to expanded r(CCUG)^exp^ repeats in DM2 but not to r(CUG)^exp^ repeats in DM1 muscle cells^75^. However, unlike MBNL family of proteins, RBFOX binding to the r(CCUG)^exp^ repeats does not result in sequestration or loss of splicing activity^75^.

We demonstrate that in DM1 patient hearts, levels of a highly active non-muscle RBFOX2_40_ splice isoform are elevated. Our data support a model where combined effects of increased CELF1 activity^19^ and reduced expression of specific miRNAs^20^ allow for selective upregulation of RBFOX2_40_ isoform in DM1 heart tissue **(Fig. 7D)**. We further propose that compared to the muscle isoform, RBFOX2_40_ stimulates the higher-order assembly of LASR complex^46, 47^, which boosts its splicing activity, resulting in altered isoform expression of proteins that can disrupt the normal cardiac rhythm and function. Indeed, we demonstrated that RBFOX2_40_ promotes the generation of pathogenic splice variants of voltage-gated sodium and potassium channels that are known to exhibit slower conduction velocity and increased susceptibility to arrhythmias^52, 54, 68, 69^. Our MD simulations showed that a RBFOX2_40_-mediated splicing switch in Na_v_1.5 transcript changes the amino acid composition of the voltage sensing element within domain I that promotes persistent lipid-interactions and stabilizes the 3_10_-helical conformation of S4 helix, which reduces the energy barrier and allows for rapid deactivation of the channel. Accordingly, we found that *tet*-inducible RBFOX2_40_ overexpression or targeted deletion of the 43-nt muscle-specific exon in *Rbfox2* gene results in prolonged PR and QT intervals, inadvertent sinus pauses, atrioventricular heart blocks, and sudden death in mice. Notably, in a recent large-scale transcriptome study of human DM1 patients, *RBFOX2* mRNA levels were inversely correlated with the strength of tibialis muscle^76^. In the future, it would be interesting to determine whether increased *RBFOX2* mRNA abundance in DM1 skeletal muscle also accompanies a corresponding splicing switch and whether the non-muscle RBFOX2_40_ isoform acts as a disease modifier in skeletal muscle as well.

The RNA toxicity hypothesis for DM1 predicts that longer CUG repeats will lead to greater sequestration of MBNL proteins, resulting in increased severity of the disease. Although the severity tends to rise with repeat length, extreme variability in penetrance of specific symptoms is observed across DM1 patient population^2^. For instance, cardiac involvement is reported in more than 80% of individuals, and conduction abnormalities can cause up to 30% of DM1 fatalities^9, 10, 13^. However, which patients have the highest risk for sudden cardiac death and what molecular events dictate these outcomes are currently unknown. Previous studies have indicated that high NKX2-5 and PKC activities can modify DM1-associated RNA toxicity in the heart^77, 78^. In our study, we noticed that RBFOX2_40_ isoform levels varied considerably among the hearts of DM1 patients. Whether the degree of RBFOX2_40_ upregulation depends on the size of CTG repeat expansion and/or if a threshold for RBFOX2_40_ levels exists that increases the susceptibility of DM1 patients to cardiac arrhythmias remain topics for future investigations. Nonetheless, the fact that normal splicing patterns for both sodium and potassium channels could be restored within days of silencing the RBFOX2_40_ expression highlights the potential of RBFOX2_40_ isoform as a therapeutic target and a biomarker for DM1 cardiac pathology.

## Supporting information

Experimental Procedures

## DATA AVAILABILITY

All raw RNA-seq data files are available for download from NCBI Gene Expression Omnibus (http://www.ncbi.nlm.nih.gov/geo/) under accession number GSE126771.

## ACKNOWLEDGEMENTS

This research was supported through NIH (R01HL126845) and Muscular Dystrophy Association (MDA514335) grants to A.K., NIH (R01HL134824; R35HL135754) to P.J.M., NIH (P41-GM104601, R01-GM122420, and R01-GM123455) to E.T., and NIH (R01AR045653 and R01HL045565) to T.A.C. C.M. was supported by American Heart Association post-doctoral fellowship (16POST29950018). S.B. was supported by the NIH Tissue microenvironment training program (T32-EB019944). All simulations were performed using the Blue Waters PRAC allocation (grant ACI1713784 to E.T.) of the National Center for Supercomputing Applications at UIUC. Three cores at UIUC supported this project: Transgenic mouse facility core, High-throughput sequencing and genotyping core and Histology-microscopy core. Cardiovascular phenotyping was carried out in part in the Biomedical Imaging Center at the Beckman Institute for Advanced Science and Technology at the University of Illinois at Urbana-Champaign and in the Mouse Cardiovascular Phenotyping Core at the Washington University School of Medicine, St. Louis, MO.

## AUTHOR CONTRIBUTIONS

C.M., S.B., and A.K. conceived the project and designed the experiments. C.M., S.B., F.L., K.L., D.J.P., S.N.K., E.R.L., N.P.M., J.H. and W.L.D. performed experiments. T.A.C. and P.J.M. provided reagents. C.M., S.B., K.L., S.N.K., E.R.L., N.P.M., E.T., P.J.M. and A.K. interpreted results and wrote the manuscript. All authors discussed the results and edited the manuscript.

