## Supplementary Figures and Tables for "Overexpression of a non-muscle RBFOX2 isoform triggers cardiac conduction defects in myotonic dystrophy"

### SUPPLEMENTARY FIGURE LEGENDS

**Supplementary Figure 1. *RBFOX2* mRNA abundance and splicing analysis in human and mouse hearts.** This supplementary figure is related to Figure 1.

**Supplementary Figure 2. miRNA regulation of *Rbfox2* in DM1 cardiac cultures.** This supplementary figure is related to Figure 2.

**Supplementary Figure 3. Effects of CELF1 overexpression on RBFOX2 protein levels in mouse heart and HL-1 cells.** This supplementary figure is related to Figure 3.

**Supplementary Figure 4. Generation of TRE-RBFOX2$_{40}$; MHCrtTA bitransgenic and *Rbfox2*$^{\Delta 43/\Delta 43}$ mice.** This supplementary figure is related to Figure 4.

**Supplementary Figure 5. Cardiac function tests of TRE-RBFOX2$_{40}$; MHCrtTA bitransgenic and *Rbfox2*$^{\Delta43/\Delta43}$ mice.** This supplementary figure is related to Figure 4.

**Supplementary Figure 6. RNA-seq analysis of cardiomyocytes isolated from Dox induced TRE-RBFOX2$_{40}$; MHCrtTA mice.** This supplementary figure is related to Figure 5.

**Supplementary Figure 7. Lipid contact frequency within the bilayer for K211 of SCN5A (6A) isoform in comparison to D211 in SCN5A (6B) isoform.** This supplementary figure is related to Figure 6.

**Supplementary Figure 8. Alternative splicing analysis of voltage-gated potassium, calcium, and sodium channels.** This supplementary figure is related to Figure 7.

### SUPPLEMENTARY TABLES

**Supplementary Table 1. Primer and oligonucleotides sequences used in qRT-PCR and cell culture experiments.** This supplemental table is related to Figure 1,2,3,4,7.

**Supplementary Table 2. Information for RNA sequencing experiments including mapping rates.** This supplemental table is related to Figure 5.

**Supplementary Table 3. Information for antibodies used in relevant Western blot experiments.** This supplemental table is related to Figure 1, 2, 3, 4, 7.

### SUPPLEMENTARY VIDEOS

**Supplementary Video 1. Interactions of D211 in SCN5A_6B with the lipid head group.** This supplemental video is related to Figure 6.

**Supplementary Video 2. Interactions of K211 in SCN5A_6A with the lipid head group.** This supplemental video is related to Figure 6.

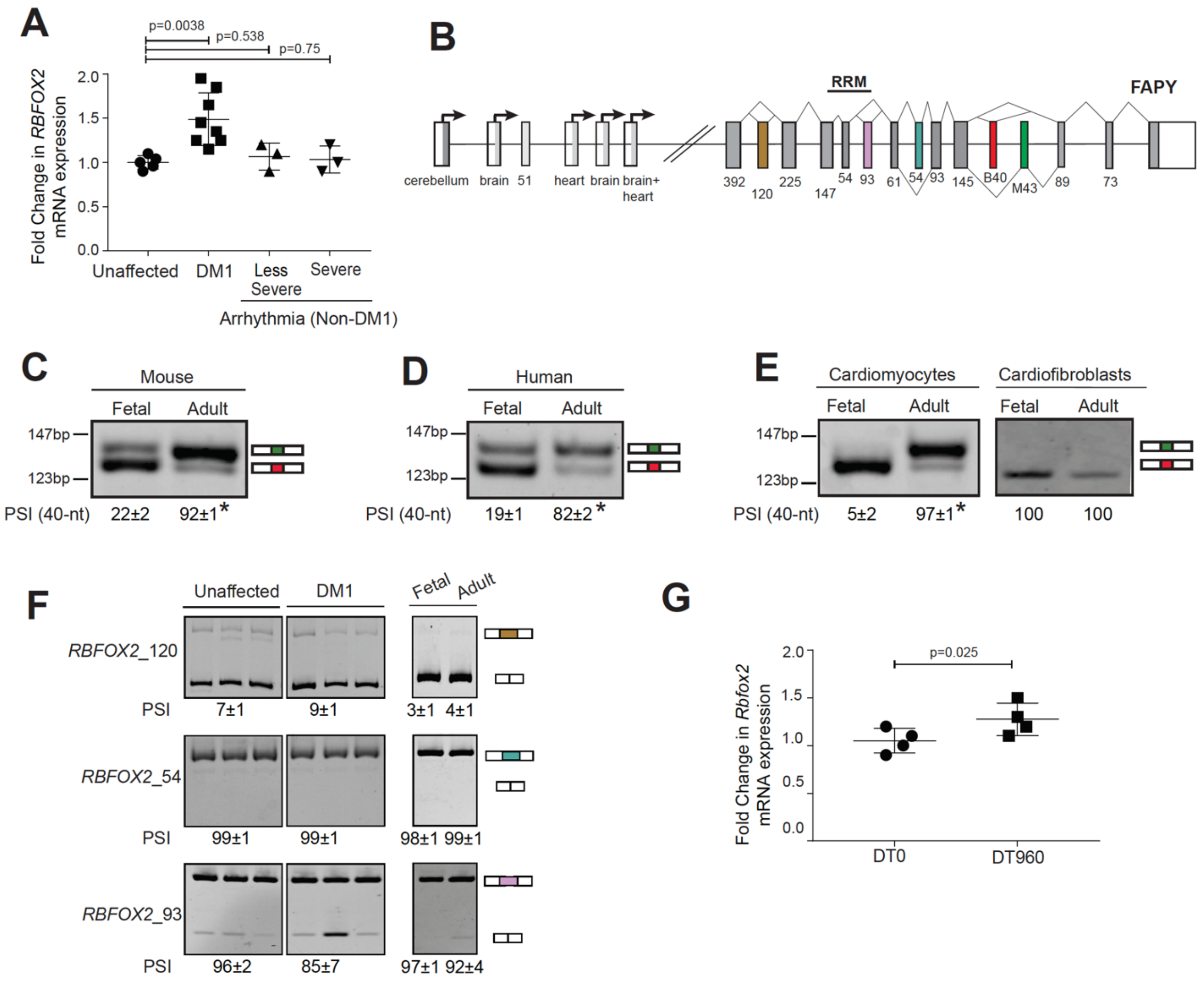


**Supplementary Figure 1. *RBFOX2* mRNA expression and splicing analysis in human and mouse hearts. (A)** Quantification of *RBFOX2* mRNA levels in unaffected (n=8), DM1 (n=9), and arrhythmic non-DM1 (Severe arrhythmia n=3, Less severe arrhythmia n=3) human heart samples. **(B)** Schematic of *Rbfox2* gene structure. Multiple promoters produce distinct first exons in specific tissues. The RNA recognition motif (RRM) is encoded by the 54-nt and 93-nt exons. The 93-nt exon in the RRM is skipped in some cases to generate a dominant negative isoform. Mutually exclusive exons M43 (muscle-expressed 43-nt exon) and B40 (brain-expressed 40-nt exon) are alternatively spliced to generate the muscle and non-muscle RBFOX2 protein isoforms. Black boxes and white boxes represent translated and untranslated sequences respectively. RT-PCR analysis monitoring the inclusion of *RBFOX2* 40-nt and 43-nt exons in fetal and adult **(C)** mice hearts (n=4), **(D)** human hearts (n=3), and **(E)** freshly isolated mouse cardiomyocytes and cardiofibroblasts (n=3). PSI: Percent Spliced In. **(F)** RT-PCR analysis monitoring the inclusion of *RBFOX2* 54-nt, 120-nt and 93-nt exons between unaffected (n=6), and DM1 (n=9) as well as fetal (n=3) and adult (n=3) human heart samples. **(G)** Quantification of *Rbfox2* mRNA levels in HL-1 cells transfected with DT0 or DT960 plasmids for 48h. All data are mean ± s.d., and p-values were derived from a parametric t-test (two-sided, unpaired), with Welch's correction. *$P$<0.05.

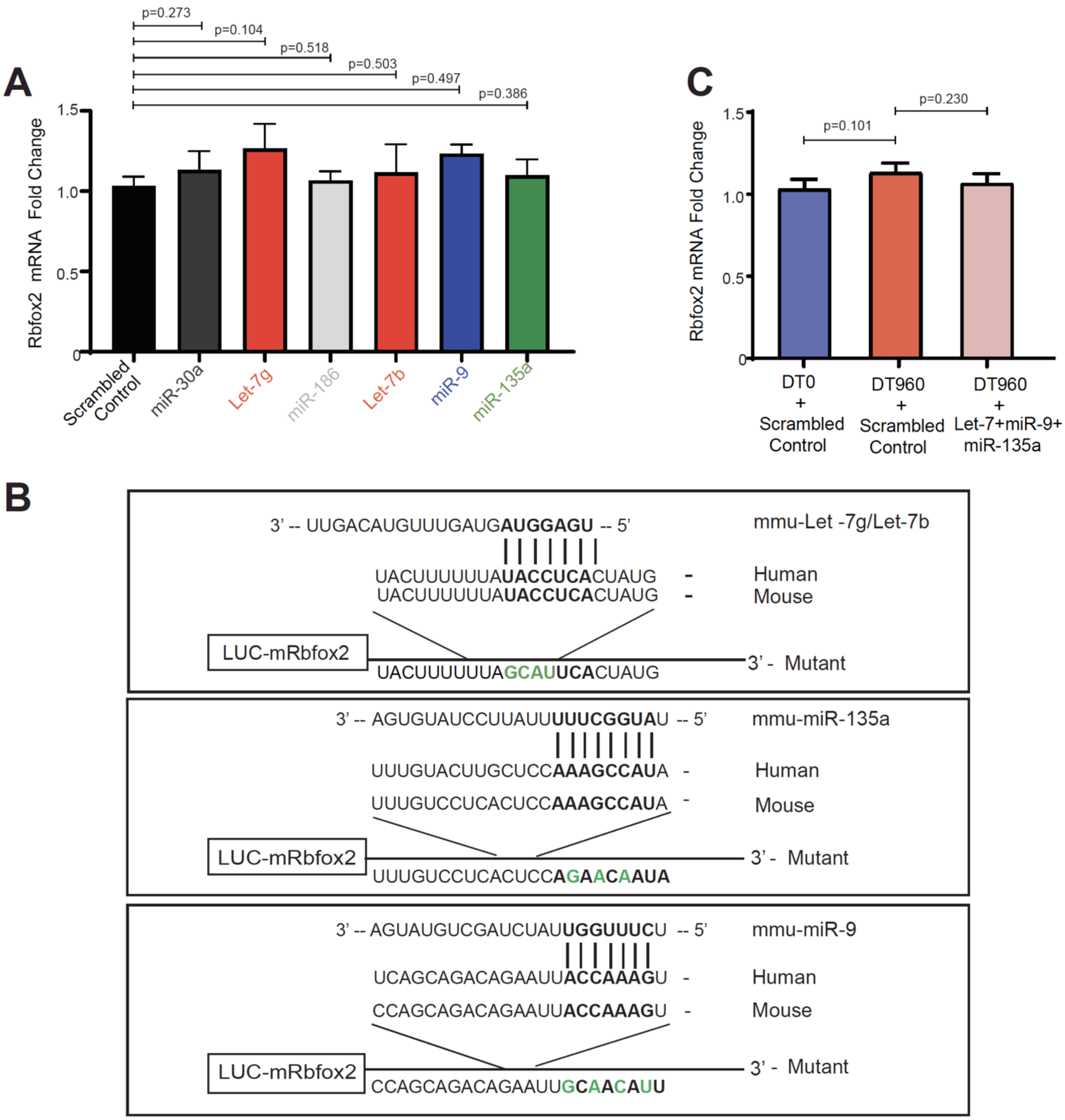


**Supplementary Figure 2. miRNA regulation of *Rbfox2* in DM1 cardiac cultures. (A)** qRT-PCR analysis of *Rbfox2* mRNA expression in HL-1 cells following treatment with scrambled control or indicated miRNA mimics. n=4 independent transfections. **(B)** Putative Let-7, miR-9 and miR-135a seed sequences in the 3′-UTRs of human and mouse *RBFOX2* transcripts. Specific mutations introduced into luciferase reporter constructs are shown in green. **(C)** qRT-PCR analysis of *Rbfox2* mRNA expression in HL-1 cells after co-transfection with DT0 or DT960 plasmids, and scrambled control or a cocktail of indicated miRNA mimics. n=4 independent transfections. All data are mean ± s.d., and p-values were derived from one-way ANOVA plus Dunnett's post-hoc test.

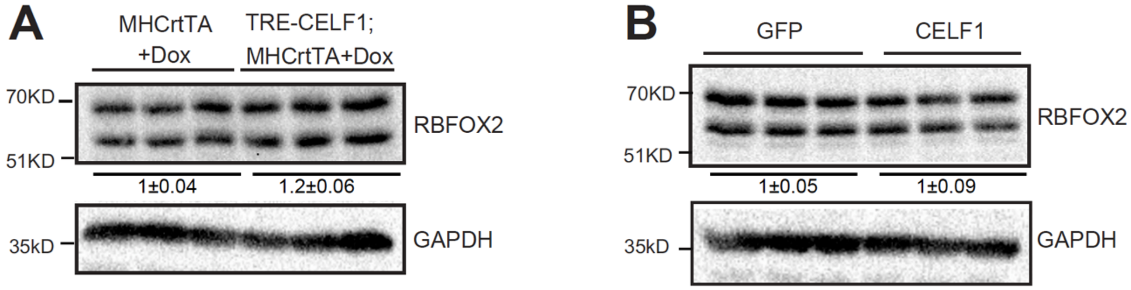


**Supplementary Figure 3. Effects of CELF1 overexpression on RBFOX2 protein levels in mouse heart and HL-1 cells.** Immunoblot analysis of RBFOX2 protein in **(A)** hearts of *tet*-inducible, heart-specific CELF1 bitransgenics (TRE-CELF1; MHCrtTA) and littermate control (MHCrtTA) mice induced with 6g/kg Dox for ten days (n=3 mice for each genotype) as well as in **(B)** HL-1 cells following infection with GFP or CELF1 expressing adenoviruses (n=3 independent transfections). All data are mean ± s.d., and p-values were derived from a parametric t-test (two-sided, unpaired), with Welch's correction.

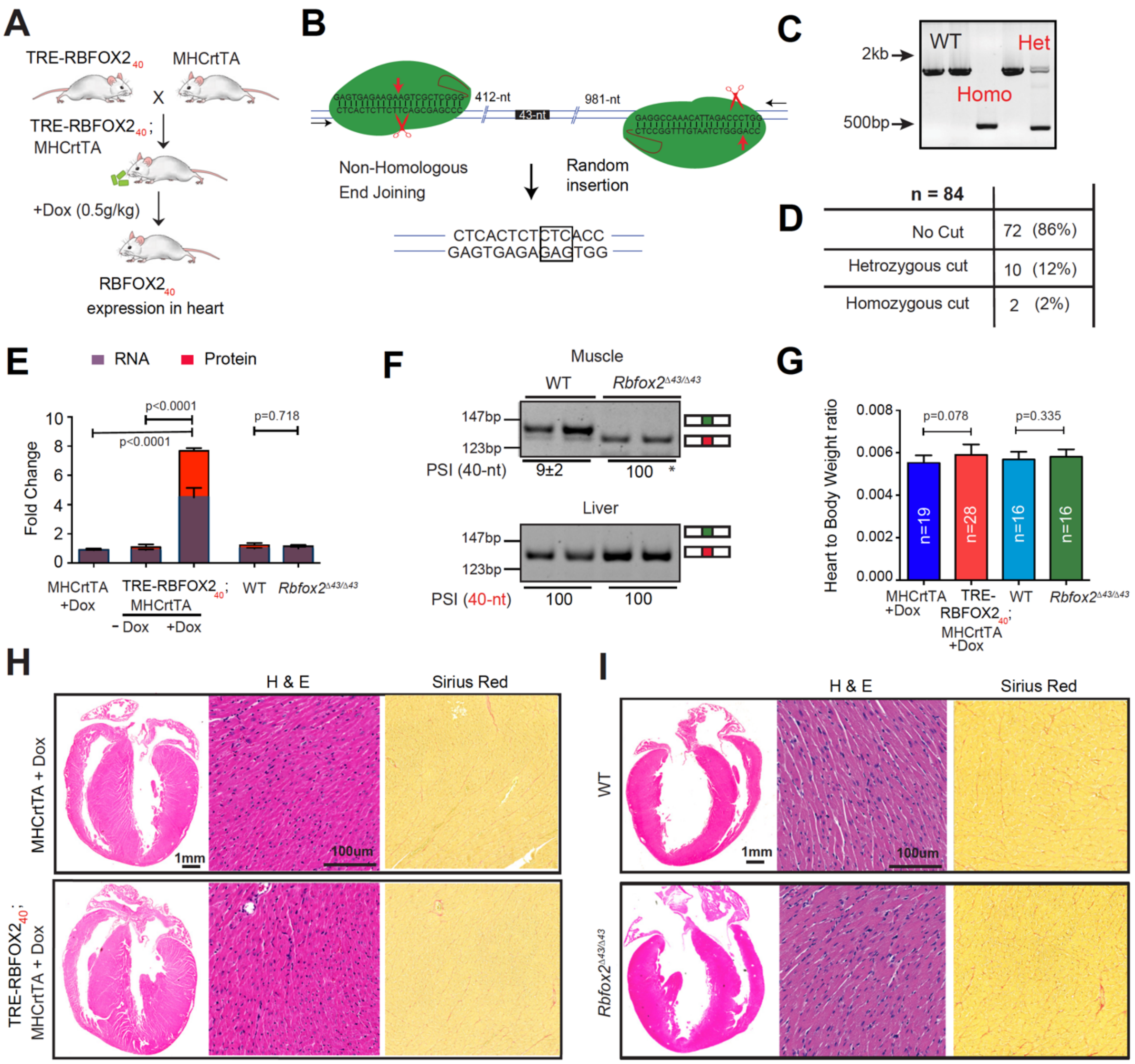


**Supplementary Figure 4. Generation of TRE-RBFOX2$_{40}$;MHCrtTA bitransgenic and *Rbfox2*$^{\Delta43/\Delta43}$ mice. (A)** Schematic of TRE-RBFOX2$_{40}$; MHCrtTA bitransgenic mice to inducibly express FLAG-tagged RBFOX2$_{40}$ protein isoform in the heart. **(B)** Schematic representation of CRISPR/Cas9 approach to generate *Rbfox2*$^{\Delta43/\Delta43}$ mice. Relevant genomic sequence with guide RNAs targeting the intronic regions flanking 43-nt exon are shown. **(C)** Genotyping results showing successful generation of *Rbfox2*$^{\Delta43/\Delta43}$ mice with 1.4kb of deletion spanning the 43-nt exon. **(D)** 12% of the screened founders were heterozygous and 2% were homozygous for the targeted deletion. **(E)** *Rbfox2* mRNA and protein expression in the hearts of hemizygous MHCrtTA, and TRE-RBFOX2$_{40}$; MHCrtTA bitransgenic mice fed 0.5g/kg Dox containing chow for 3 days, as well as of wildtype (WT) and *Rbfox2*$^{\Delta43/\Delta43}$ mice. n=5 for each genotype. **(F)** RT-PCR analysis of *Rbfox2* 43-nt and 40-nt exons in the muscle and liver tissues from WT and *Rbfox2*$^{\Delta43/\Delta43}$ mice. PSI: Percent Spliced In; n=4-6 mice for each genotype. **(G)** Heart-to-body weight ratios, Representative **(H)** H&E, and **(I)** Sirius Red staining of hearts from indicated genotypes. All data are mean ± s.d., and p-values were derived from a parametric t-test (two-sided, unpaired), with Welch's correction.

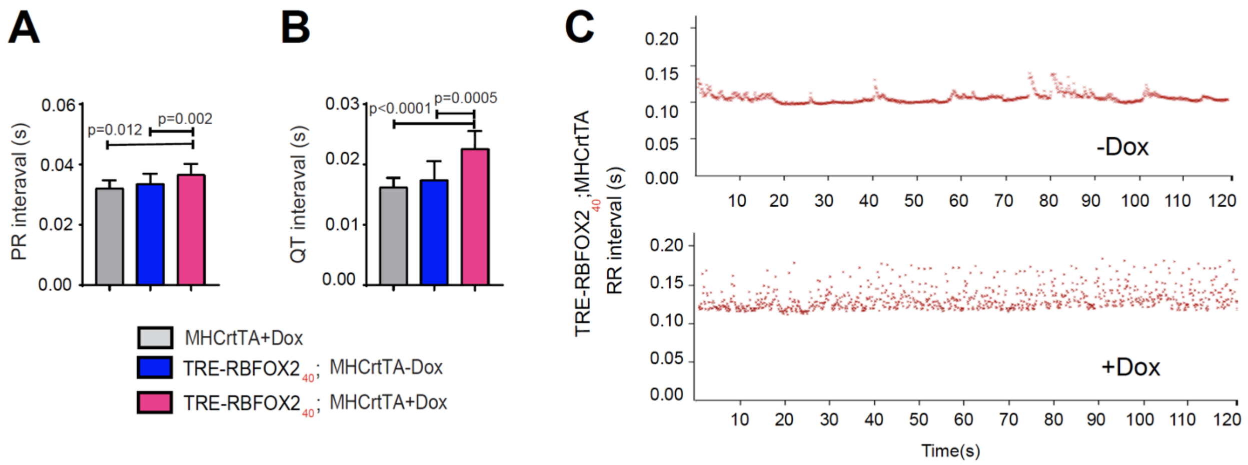


**D**

| Parameter | MHCrtTA +Dox | TRE-RBFOX2$_{40}$; MHCrtTA+Dox | WT | *Rbfox2*$^{\Delta43/\Delta43}$ |
|---|---|---|---|---|
| Heart Rate(BPM) | 360.21 | 336.84 | 358.81 | 365.5 |
| Diameter;s (mm) | 2.78 | 3.05 | 1.72 | 2.00 |
| Diameter;d (mm) | 4.19 | 3.97 | 3.38 | 3.50 |
| Volume;s (ul) | 29.43 | 37.13 | 10.15 | 14.42 |
| Volume;d (ul) | 78.78 | 69.71 | 47.5 | 51.4 |
| Stroke Volume (ul) | 49.34 | 32.57 * | 37.34 | 36.9 |
| Ejection Fraction (%) | 62.44 | 46.15 * | 80.42 | 73.61 |
| Fractional Shortening (%) | 33.51 | 23.01 * | 49.81 | 43.24 |
| Cardiac Output (mL/min) | 17.71 | 11.44 | 13.34 | 13.69 |

**Supplementary Figure 5. Cardiac function tests of TRE-RBFOX2$_{40}$; MHCrtTA bitransgenic and *Rbfox2*$^{\Delta43/\Delta43}$ mice.** Surface ECG analysis showing **(A)** PR, and **(B)** QT intervals of MHCrtTA mice (n=10) fed 0.5g/kg Dox containing chow for 9 days, and the TRE-RBFOX2$_{40}$; MHCrtTA bitransgenic mice (n=14) 48h before and 9 days after 0.5g/kg Dox administration. **(C)** Variation in RR interval from ECG analysis in TRE-RBFOX2$_{40}$; MHCrtTA bitransgenic mice 48h before and 9 days after 0.5g/kg Dox administration (n=4). **(D)** Echocardiographic analysis of MHCrtTA (n=10), and TRE-RBFOX2$_{40}$; MHCrtTA bitransgenics induced with 0.5g/kg Dox for 9 days (n=14), as well as wildtype (WT, n=6) and *Rbfox2*$^{\Delta43/\Delta43}$ (n=8) mice. All data are mean ± s.d., and p-values were derived from a parametric t-test (two-sided, unpaired for **A, B, D** and paired for **C**), with Welch's correction.

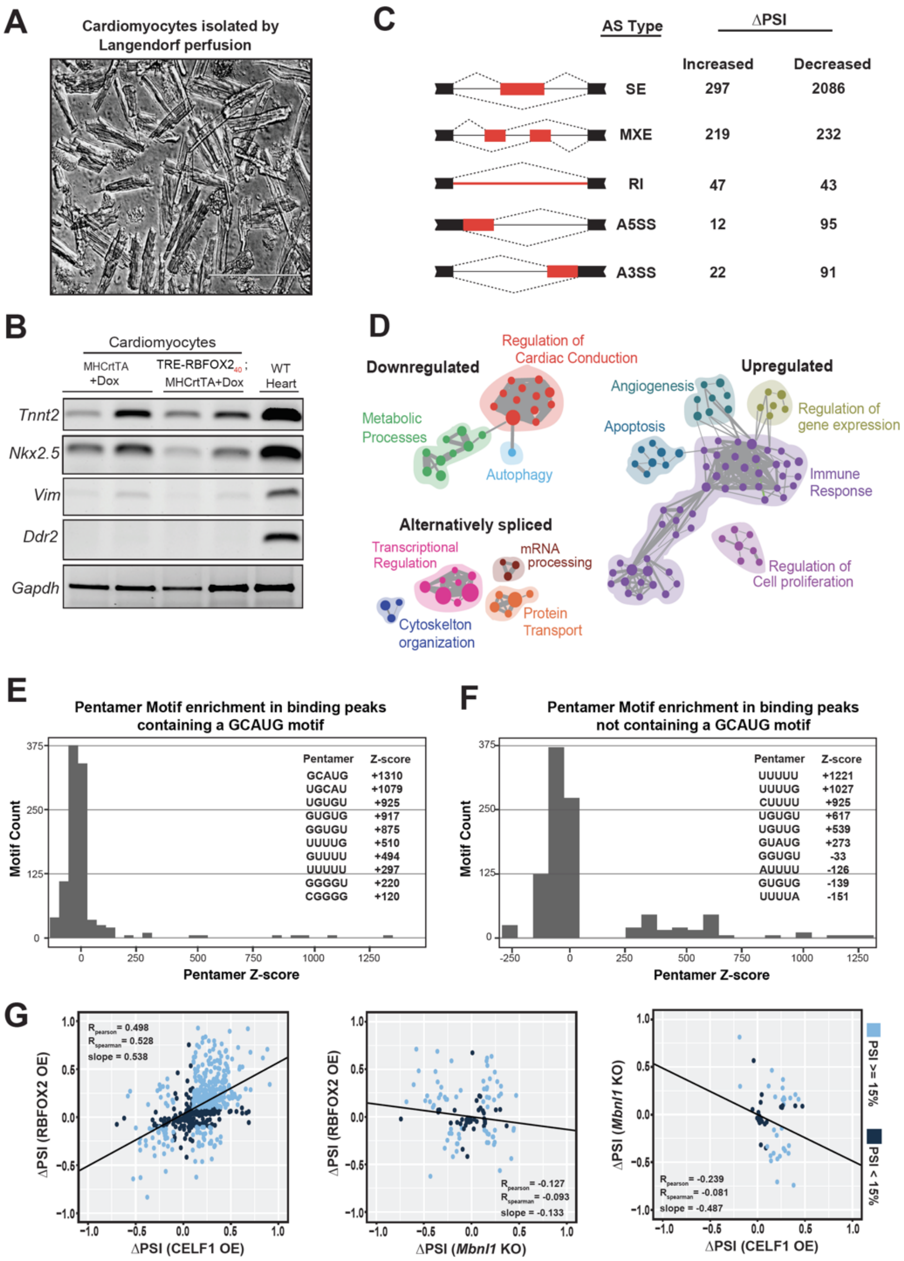


**Supplementary Figure 6. RNA-seq analysis of cardiomyocytes isolated from Dox induced TRE-RBFOX2$_{40}$; MHCrtTA mice. (A)** Photograph of isolated adult ventricular cardiomyocytes from TRE-RBFOX2$_{40}$; MHCrtTA bitransgenics induced with 0.5g/kg Dox for 3 days. **(B)** RT-PCR analysis of cardiomyocyte (*Tnnt2*, *Nkx2.5*) and cardiofibroblast (*Vim*, *Ddr2*) markers in the indicated samples. **(C)** Classification of splicing event types that change significantly after RBFOX2$_{40}$ overexpression. ΔPSI: Difference in Percent Spliced In. **(D)** Gene ontology analysis of differentially expressed and spliced genes following RBFOX2$_{40}$ overexpression. Distribution of z-scores of 1024 pentameric motifs in sequences **(E)** with and **(F)** without GCAUG motif near RBFOX2$_{40}$-regulated exons in cardiomyocytes. Top 10 enriched motifs with relevant z-scores are listed. **(G)** Scatter plots of pairwise comparisons for ΔPSIs following RBFOX2$_{40}$ overexpression (OE) in cardiomyocytes, and CELF1 overexpression and *Mbnl1*$^{\Delta E3/\Delta E3}$ in mouse hearts.

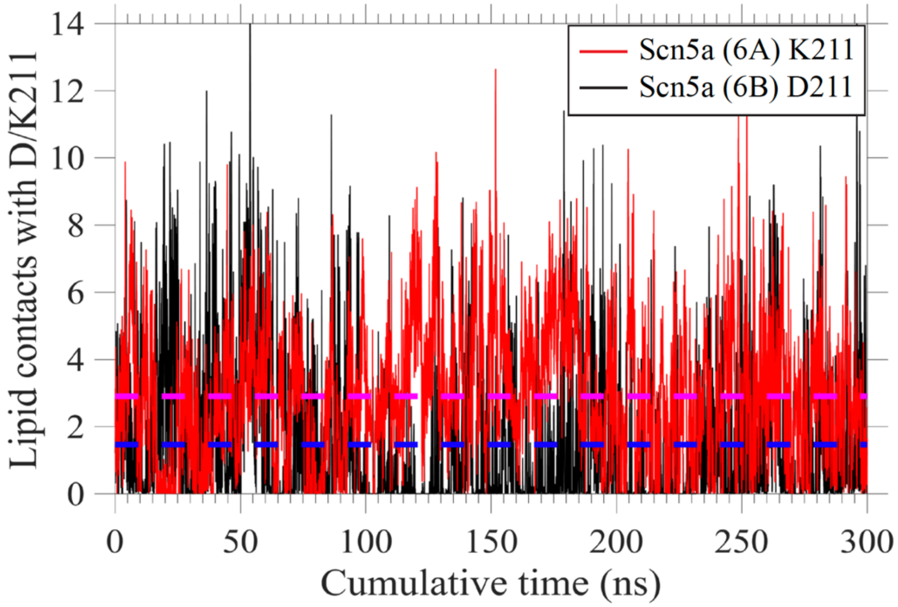


**Supplementary Figure 7. Lipid contact frequency within the bilayer for K211 of SCN5A (6A) isoform in comparison to D211 in SCN5A (6B) isoform.** Lipid contacts were calculated between heavy atoms of lipid molecules and the sidechain carboxylate (Asp) or ammonium (Lys) groups. The magenta and blue horizontal lines show the average lipid contacts for SCN5A (6A) isoform and SCN5A (6B) isoform respectively. All three independent simulations for each isoform were concatenated into one single trajectory for the analysis. The average contact counts between K211 and lipid molecules was 2.9 which was double than that of D211.

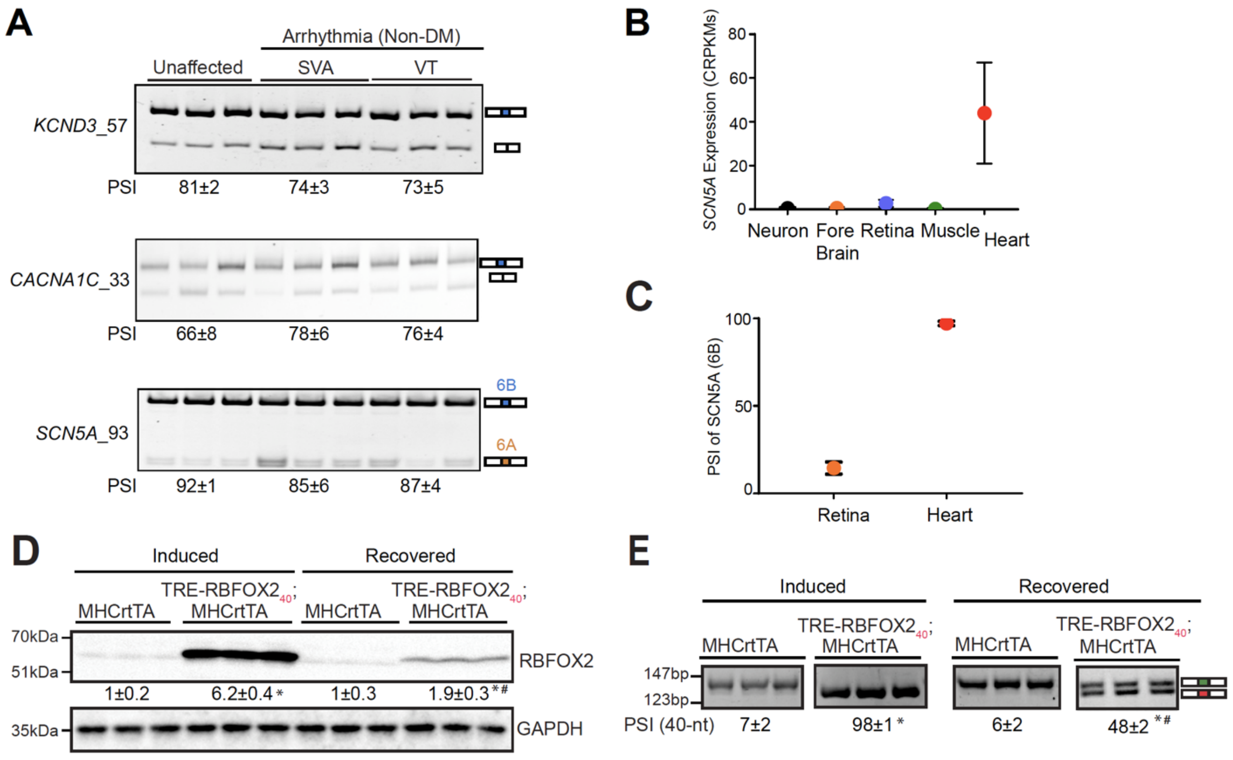


**Supplementary Figure 8. Alternative splicing analysis of voltage-gated potassium, calcium, and sodium channels. (A)** RT-PCR analysis comparing the inclusion of 57-nt exon in *KCND3*, 33-nt exon in *CACNA1C*, and 93-nt mutually exclusive exons (6A and 6B) in *SCN5A* transcripts between unaffected (n=3), and arrhythmic non-DM [SVA: Sustained Ventricular Arrhythmia (n=3); and VT: Ventricular Tachycardia (n=3)] human heart samples. Percent Spliced In (PSI) values are shown below the gel images. **(B)** Comparison of *SCN5A* expression in different tissues from VastdB database. **(C)** Adult *SCN5A* (6B) isoform expression in human retina vs. heart from VastdB database. **(D)** Immunoblot analysis of RBFOX2 protein in the hearts of TRE-RBFOX2$_{40}$; MHCrtTA and littermate control (MHCrtTA) mice induced with 0.5g/kg Dox for 24h followed by a 3-day recovery period. Quantification of relative band intensities for RBFOX2 normalized to GAPDH are shown below the gel image. n=3 mice for each genotype/condition. **(E)**. RT-PCR analysis of *Rbfox2* 43-nt and 40-nt exons in the hearts of TRE-RBFOX2$_{40}$; MHCrtTA and littermate control (MHCrtTA) mice induced with 0.5g/kg Dox for 24h followed by a 3-day recovery period. PSI values are shown below the gel images. n=3 mice for each genotype/condition. All data are mean ± s.d., and p-values were derived from a parametric t-test (two-sided, unpaired), with Welch's correction for **A**, and from one-way ANOVA plus Dunnett's post-hoc test for **D** and **E**. *P<0.05 (between MHCrtTA and TRE-RBFOX2$_{40}$; MHCrtTA), and $^{\#}$P<0.05 (between TRE-RBFOX2$_{40}$; MHCrtTA (induced) and TRE-RBFOX2$_{40}$; MHCrtTA (recovered).

### SUPPLEMENTARY TABLES

**Supplementary Table 1. Primer and oligonucleotides sequences used in relevant genotyping, qRT-PCR, RT-PCR and cell culture experiments.** This supplemental table is related to Figure 1,2,3,4,7.

| Target Gene | Type | Primer Sense | Sequence (5'->3') |
|---|---|---|---|
| *MHCrtTA* | Genotype | F | CTGGGTTGCGTGTTGGAAGATC |
| *MHCrtTA* | Genotype | R | GTGGGAGATCGAGCAGGCCCTCG |
| 3'TRE-*Rbfox2* | Genotype | F | ACAGCCTGCTACTGCAACC |
| 3'TRE-*Rbfox2* | Genotype | R | GCGATGCAATTTCCTCATTT |
| 5'TRE-*Rbfox2* | Genotype | F | AAGTGAAAGTCGAGCTCGGTA |
| 5'TRE-*Rbfox2* | Genotype | R | GTTGTTGTTGGCTCCTGGTT |
| *Rbfox2*$^{\Delta 43/\Delta 43}$ | Genotype | F | CAAGACACCTTCTTTCTACCTG |
| *Rbfox2*$^{\Delta 43/\Delta 43}$ | Genotype | R | GCTGGAGCTGTTAAACTGATG |
| *Rbfox2*$^{\Delta 43/\Delta 43}$_43 | Genotype | F | ATCCATCACCATGCCTTTGC |
| *Rbfox2*$^{\Delta 43/\Delta 43}$_43 | Genotype | R | GGAGCTGGAATGGTTAGTAT |
| *RBFOX1*_43_40_human | RT-PCR | F | GCACCGTGTACAACACCTTC |
| *RBFOX1*_43_40_human | RT-PCR | R | ACTGTAGGCAGCGGCAGT |
| *RBFOX2*_43_40_human | RT-PCR | F | GGTACCTCCAACAGCCATCC |
| *RBFOX2*_43_40_human | RT-PCR | R | GTGTACACCCTGCCATAA |
| *Rbfox2*_43_40_mouse | RT-PCR | F | GGTACCTCCAACAGCCATCC |
| *Rbfox2*_43_40_mouse | RT-PCR | R | GTGTACACCCTGCCGTAA |
| *RBFOX2*_54_human | RT-PCR | F | CCCTGTCTGCATCAGCACTA |
| *RBFOX2*_54_human | RT-PCR | R | CAGAAGGTGGAGCACAGACA |
| *RBFOX2*_120_human | RT-PCR | F | GCCGGCATAGTCTTGAGTGT |
| *RBFOX2*_120_human | RT-PCR | R | TTCCTTCAGCCATCTGCCTG |
| *RBFOX2*_93_human | RT-PCR | F | CAGTTTGGCAAAATCCTAGATG |
| *RBFOX2*_93_human | RT-PCR | R | GGTGTGACCATCTTCTTATTGG |
| *CACNA1C*_33_ human | RT-PCR | F | AAATCGCCATGAACATCCTC |
| *CACNA1C* _33_ human | RT-PCR | R | TTGATGAAGGTCCACAGCAG |
| *Cacna1c* _33_ mouse | RT-PCR | F | GAGCTGCCTCCTCAAAATCG |
| *Cacna1c* _33_ mouse | RT-PCR | R | AAGAGGCGGAAGAAGGTGAT |
| *KCND3*_ 57_ human | RT-PCR | F | CCAGAAGAGGAGCACATGGG |
| *KCND3*_ 57_ human | RT-PCR | R | GGGACTTCTTGTGGATGGGT |
| *Kcnd3*_ 57_ mouse | RT-PCR | F | GGCAAGACCACCTCACTCAT |
| *Kcnd3*_ 57_ mouse | RT-PCR | R | TGGCTGGACAGAGAAGGACT |
| *SCN5A*_93_ human | RT-PCR | F | CTTCTGCCTGCACGCGTTCAC |
| *SCN5A*_93_ human | RT-PCR | R | CAGAAGACTGTGAGGACCATC |
| *Scn5a*_93_ mouse | RT-PCR | F | CTTCTGCCTGCATGCGTTCAC |
| *Scn5a*_93_ mouse | RT-PCR | R | CAGAAGACAGTAAGGACCATC |

| | | | |
|---|---|---|---|
| *Tnnt2* | RT-PCR | F | CGGAAGAGTGGGAAGAGACA |
| *Tnnt2* | RT-PCR | R | TTCCCACGAGTTTTGGAGAC |
| *Nkx2.5* | RT-PCR | F | AAGCAACAGCGGTACCTGTC |
| *Nkx2.5* | RT-PCR | R | GGGTAGGCGTTGTAGCCATA |
| *Vim* | RT-PCR | F | TGAAGGAAGAGATGGCTCGT |
| *Vim* | RT-PCR | R | TTGAGTGGGTGTCAACCAGA |
| *Ddr2* | RT-PCR | F | CAAGATCATGTCTCGGCTCA |
| *Ddr2* | RT-PCR | R | GCCCTGGATCCGGTAGTAAT |
| *Rbfox2*_mouse | qRT-PCR | F | GGGAAGCCAGGAACTAAAGG |
| *Rbfox2*_mouse | qRT-PCR | R | TGTGTCCCTAGGCAATGATG |
| *RBFOX2*_human | qRT-PCR | F | GGTACCTCCAACAGCCATCC |
| *RBFOX2*_human | qRT-PCR | R | GTGTACACCCTGCCATAA |
| *Celf1* siRNA | | | CGUUUGGACAGAUUGAAGAtt |
| *Rbfox2*_3'-UTR | PCR | F | GACTTGAATTCAAGGCCTCAGTGACG TGAGACCCCTGCAAATGGG |
| *Rbfox2*_3'-UTR | PCR | R | CGACTCACTATAGTTCTAGATGGTGT TTCTCTCTTTTATTTAAAAAC |
| Mut Let-7 | SDM | F | TTCAAAGAACATAGTGAATGCTAAA AAAGTATTCTCTCTTTTTTGTTTGTT TTCCTCTTC |
| Mut Let-7 | SDM | R | GAAGAGGAAAACAAACAAAAAAGAG AGAATACTTTTTTAGCATTCACTATG TTCTTTGAA |
| Mut miR-9 | SDM | F | AGCCATCAGCACCAGATCCAATGTTG CAATTCTGTCTGCTGGCTTCTGATC |
| Mut miR-9 | SDM | R | GATCAGAAGCCAGCAGACAGAATTG CAACATTGGATCTGGTGCTGATGGCT |
| Mut miR-135a | SDM | F | CTGGAAGATGGAAATTCCTATTGTT CTGGAGTGAGGACAAACTGCCT |
| Mut miR-135a | SDM | R | AGGCAGTTTGTCCTCACTCCAGAACA ATAGGAATTTCCATCTTCCAG |

SDM: site directed mutagenesis.

**Supplementary Table 2. Information for RNA sequencing experiments including mapping rates for individual files.** This supplemental table is related to Figure 5.

| Sample Name | SRA File ID | Read Length | Total Reads | Mapped Reads | % Mapped |
|---|---|---|---|---|---|
| MHCrtTA 3 day Dox Replicate 1 | This paper | 2X100 | 99501365 | 71572733 | 71.93 |
| MHCrtTA 3 day Dox Replicate 2 | This paper | 2X100 | 90376372 | 65357985 | 72.32 |
| TRE-RBFOX2$_{40}$; MHCrtTA 3 day Dox Replicate 1 | This paper | 2X100 | 97577814 | 67507903 | 69.18 |
| TRE-RBFOX2$_{40}$; MHCrtTA 3 day Dox Replicate 2 | This paper | 2X100 | 90249175 | 60344770 | 66.86 |
| Human Control Replicate 1 | SRR1971626 | 2X100 | 84000000 | 78961429 | 94 |
| Human Control Replicate 2 | SRR1971627 | 2X100 | 170806642 | 160968260 | 94.24 |
| Human Control Replicate 3 | SRR1971628 | 2X100 | 202795863 | 190446117 | 93.91 |
| Human DM1 Replicate 1 | SRR1971629 | 2X100 | 189993197 | 179173777 | 94.31 |
| Human DM2 Replicate 2 | SRR1971630 | 2X100 | 157762296 | 149276370 | 94.62 |
| Human DM3 Replicate 3 | SRR1971631 | 2X100 | 189593674 | 179657888 | 94.76 |
| *Mbnl1* WT Replicate 1 | SRR533627 | 1X50 | 13107864 | 8358911 | 63.77 |
| *Mbnl1* WT Replicate 2 | SRR533628-29 | 1X50 | 29500486 | 17858551 | 60.54 |
| *Mbnl1* WT Replicate 3 | SRR533630-31 | 1X50 | 27981848 | 16870533 | 60.29 |
| *Mbnl1* WT Replicate 4 | SRR533632 | 1X50 | 13563047 | 8110656 | 59.8 |
| *Mbnl1* WT Replicate 5 | SRR533633 | 1X50 | 9681738 | 5694707 | 58.82 |
| *Mbnl1*$^{\Delta E3/\Delta E3}$ Replicate 1 | SRR533622 | 1X50 | 14242938 | 8431840 | 59.2 |
| *Mbnl1*$^{\Delta E3/\Delta E3}$ Replicate 2 | SRR533623 | 1X50 | 13159419 | 7562838 | 57.47 |
| *Mbnl1*$^{\Delta E3/\Delta E3}$ Replicate 3 | SRR533624 | 1X50 | 13379187 | 7841675 | 58.61 |
| *Mbnl1*$^{\Delta E3/\Delta E3}$ Replicate 4 | SRR533625 | 1X50 | 10092421 | 5861325 | 58.08 |
| *Mbnl1*$^{\Delta E3/\Delta E3}$ Replicate 5 | SRR533626 | 1X50 | 16224009 | 9088998 | 56.02 |
| TRE-CELF1; MHCrtTA 3 day Replicate 1 | SRR1205709 | 2X35 | 11583447 | 9619626 | 83.05 |
| TRE-CELF1; MHCrtTA 3 day Replicate 2 | SRR1205711 | 2X35 | 28294445 | 22464788 | 79.4 |
| TRE-CELF1; MHCrtTA 3 day Replicate 3 | SRR1205712-13 | 2X35 | 25072150 | 12137386 | 48.41 |
| MHCrtTA 3 day Dox Replicate 1 | SRR1205699 | 2X35 | 13888713 | 11462967 | 82.53 |
| MHCrtTA 3 day Dox Replicate 2 | SRR1205700 | 2X35 | 12478885 | 10107181 | 80.99 |
| MHCrtTA 3 day Dox Replicate 3 | SRR1205701 | 2X35 | 11335507 | 9439782 | 83.28 |

**Supplementary Table 3. Information for antibodies used in relevant Western blot and Immunohistochemistry experiments.** This supplemental table is related to Figure 1, 2, 3, 4, 7.

| Target | Type | Assay | Origin | Dilution | Source | Catalogue No. |
|---|---|---|---|---|---|---|
| RBFOX2 | $1^0$ | WB | Rabbit | 1:1000 | Bethyl | A300-864A |
| CELF1 | $1^0$ | WB | Mouse | 1:1000 | Santa Cruz | sc-20003 |
| GAPDH | $1^0$ | WB | Rabbit | 1:10000 | Sigma-Aldrich | G9545-200UL |
| Anti-Mouse IgG | $2^0$ | WB | Goat | 1:5000 | Bio-Rad | 1721011 |
| Anti-Rabbit IgG | $2^0$ | WB | Goat | 1:5000 | Thermo Fisher | SA5-10033 |
