## Supplementary material for "Overexpression of a non-muscle RBFOX2 isoform triggers cardiac conduction defects in myotonic dystrophy": Experimental Procedures

### Animal models and human samples

Use and care of laboratory animals were followed according to the National Institutes of Health (NIH) and guidelines set by the Institutional Animal Care and Use Committee (IACUC) at University of Illinois, Urbana-Champaign (UIUC). Animals used in the study were identified using ear tags and genotyped before weaning age. Experiments within the study were not gender-specific and specimens included both male and female animals. Whole heart tissues and cardiomyocytes were isolated from mice following the IACUC guidelines for euthanasia and anesthesia.

For generating *Rbfox2*$_{40}$ non-muscle isoform overexpression mouse model, the mouse RBFOX2$_{40}$ cDNA, harboring a N-terminus sequence coding for FLAG-tag, was expressed from a transgene with a TRE/minimal CMV promoter, *RBFOX2* ORF containing non-muscle 40-nt (B40) exon and bovine growth hormone polyadenylation site and 3' flanking genomic segment for proper mRNA 3'-end formation. The linearized transgene construct was subjected to pronuclear injection in the transgenic mouse facility at UIUC, using standard methods to generate TRE-RBFOX2$_{40}$ transgenic mice that were maintained on a C57BL/6J background. MHCrtTA transgenic mice (FVB/N-Tg (Myh6rtTA)1Jam) expressing a codon-optimized rtTA variant specifically in cardiomyocytes were commercially obtained (RRID: MMRRC_010478)[1]. Henceforth, mice reported were the F1 progeny of TRE-RBFOX2$_{40}$ and MHCrtTA mating and were, therefore, hemizygous for one or both transgenes. Primers used to genotype both transgenes are listed in **Supplementary Table 1**. RBFOX2$_{40}$ isoform expression in 8- to 12-week-old bitransgenic animals was induced through doxycycline (Dox) in the food (0.5g Dox/kg food, Harlan, KY).

To delete the muscle-specific 43-nt exon in *Rbfox2*, two single guide RNAs (sgRNAs) were purchased (Sigma-Aldrich, St. Louis, Missouri) that flank the genomic region across the

43-nt exon of *Rbfox2* (5'sgRNA: 5'GAGTGAGAAGAAGTCGCTCGGG3' and 3'sgRNA: 5'GAGGCCAAACATTAGACCCTGG3'). After complex formation with clustered regularly interspaced short palindromic repeat (CRISPR)-associated protein 9 (Cas9) (CP01-20, Newbury Park, California), the Cas9-sgRNA mixture was delivered into the cytoplasm of ~100 pronuclear stage zygotes (C57BL/6J). Injected zygotes were transferred into pseudo-pregnant ICR females (25-30 zygotes/female). The sgRNAs produced double-stranded cuts, which removed the targeted *Rbfox2* genomic sequence **(Supplementary Figure 4)**. To identify the positive founder animals wherein, double-stranded breaks induced by Cas9 resulted in non-homologous end joining (NHEJ) repair and created a null allele, mice were genotyped using standard PCR on tail-clip derived genomic DNA. Primers flanking the 2 sgRNA sites were designed **(Supplementary Table 1)** to amplify a smaller deletion amplicon compared with the WT amplicon. An additional primer pair was designed within the *Rbfox2* 43-nt exon to confirm deletion by the absence of the amplicon in PCR reactions **(Supplementary Table 1)**. The germline deletion of 43-nt exon was further confirmed by sanger sequencing of the F1 progeny mice homozygous for the 43-nt deletion (*Rbfox2*$^{\Delta 43/\Delta 43}$).

Human fetal (22-week old) heart RNAs were purchased from Clonetech Laboratories, Inc., (Mountain View, CA). Three unaffected human adult heart RNAs were from 51-year-old Caucasian male were purchased from Clonetech Laboratories, Inc., (Mountain View, CA), pooled from 3 male Caucasians, ages: 30-39 purchased from Takara Bio USA, Inc., (Mountain View, CA) and 24-year-old male RNA from Bio chain Institute Inc., (Newark, CA). Human DM1 RNA samples and other unaffected samples were from a 50-year-old male (respiratory failure [RF]), 48-year-old female (1,500 repeats, RF), 55-year-old male (pneumonia [PN]), 52-year-old female (>1,000 repeats, RF), 46-year-old male (PN), 50-year-old female (RF), 53-year-old male (unknown cause), 26-year-old male (glioma), 55-year-old male (pulmonary embolism). Unaffected heart samples were pooled autopsy samples ranging from 21- to 55-year-old individuals. Human non-DM cardiac arrhythmia samples (severe) were from 26-year-old male (Ventricular septal defects since infancy, sudden death), 47-year-old male (non-ischemic cardiomyopathy, >20 Ejection fraction [EF]), 56-year-old male (Sinus tachycardia, left posterior fascicular block, abnormal ECG). All three severe arrhythmic patients suffered from sudden cardiac arrest. Three less severe non-DM arrhythmia patient samples had a history of Ventricular tachycardia but ECG within normal limits at the time of death. All individuals in the less severe group also suffered from ischemic heart failure. They were from 50-year-old male (>15 EF), 56-year-old female (>15 EF), 64-year-old male (>24 EF).

**Conscious telemetry recordings and Surface ECGs**

Telemetry studies were performed in Mouse Cardiovascular Phenotyping Core, Washington University School of Medicine in St. Louis, with 3-month old TRE-RBFOX2$_{40}$; MHCrtTA (n=2 male, n=2 female) mice. ETA-F10 implantable radio frequency transmitters for ECG (Data Sciences International Inc.) were implanted subcutaneously in the posterior neck of adult mice. Leads were tunneled to the anterior chest in lead II position. After a post-implant recovery period of one week, ambulatory heart rhythm and heart rate (HR) were monitored in unrestrained, caged mice. After baseline recording for 24h, 0.1g/kg Dox containing diet was administered to induce expression of the RBFOX2$_{40}$ transgene. Recordings were obtained continuously at 1 kHz for nine days. The ECG Analysis module for Lab Chart (Data Sciences International Inc.) was used to automatically quantify heart rate, PR, RR, QRS, and QT intervals and the voltages of ECG recordings. Lab Chart was used to manually identify ECG events, including, premature ventricular contractions (PVCs), sinus pause (lasting at least 1.5-times as long as the preceding RR interval) and atrioventricular conduction block. Surface ECGs were captured from leads I, II and III with a MP150 data acquisition system (BIOPAC Systems Inc., Goleta, California). Two to three month-old wild type C57BL/6J (n=6 males, n=4 females ), *Rbfox2*$^{\Delta 43/\Delta 43}$ (n=8 female, n=6 female), MHCrtTA with Dox (n=3 male, n=2 female),

TRE-RBFOX2$_{40}$; MHCrtTA with and without Dox (n=6 male, n=4 female) were anesthetized in an induction chamber using 2.5% isoflurane in 100% oxygen and maintained during data collection with 1.5% isoflurane. Body temperature was monitored by a rectal probe. Data were collected for 2 minutes per mouse and annotated for analysis using ACQ Knowledge software (BIOPAC Systems Inc., Goleta, California).

**Echocardiography**

*In vivo* cardiac function and morphology were evaluated using a Vevo 2100 ultrasound machine equipped with a 40 MHz transducer-MS550S (VisualSonics, Toronto, Ontario, Canada) at the Beckman Institute for Advanced Science and Technology (UIUC). Mice were first anesthetized in an induction chamber using 2.0% isoflurane in 100% oxygen and then the anesthesia (1% Isoflurane) maintained by nose cone delivery. During the imaging procedure, mice were transferred to a heated ECG platform for heart rate monitoring. Body temperature was maintained at 37°C, monitored through a rectal probe. Two-dimensional M-mode echocardiography images were taken in the short-axis position for each animal. Data analysis was performed using the VisualSonics Vevo Lab analysis package. Three M-mode tracings were analyzed and averaged for every animal.

**Histology**

Heart tissues from Dox-induced MHCrtTA and TRE-RBFOX2$_{40}$; MHCrtTA, as well as wildtype and *Rbfox2*$^{\Delta 43/\Delta 43}$ mice were harvested and fixed overnight in 10% neutral-buffered formalin, embedded in paraffin, and sectioned (5 µm thickness). Unstained slides were deparaffinized in xylene (two treatments, 5 min each), rehydrated sequentially in ethanol (2 min each in 100%, 95%, and 80%), and washed for 3 min in water. For Hematoxylin and eosin (H&E) staining, sections were washed in Hematoxylin (2 min) and Eosin (1 min) successively, cover slipped with Permount medium, and imaged on Hamamatsu Nanozoomer. For Sirius

Red staining, sectioned tissues were stained in picro-sirius red for an hour, washed in acidified water, successively dehydrated in 100% ethanol, finally cleared in xylene and mounted.

**Isolation of neonatal and adult cardiomyocytes**

Neonatal cardiomyocytes were isolated from wildtype C57BL/6J mice with a neonatal mouse cardiomyocyte isolation kit (Cellutron Life Tech Highland Park, NJ, USA; nc-6031) as previously described[2]. Cells from 12 to 18 hearts were pooled, pre-plated for 2h on an uncoated dish to separate fibroblasts from cardiomyocytes. Adult cardiomyocytes from MHCrtTA and TRE-RBFOX2$_{40}$; MHCrtTA mice—induced with 0.5g/kg Dox containing diet for 3 days—were isolated by cannulating the hearts through the aorta and perfusing on a Langendorff apparatus  for 4 min at 3 ml/min with perfusion buffer (NaCl 58.4mM, KCl 74.55mM, MgSO4 120.4mM, Na2HPO4 142mM, KH2PO4 136.1mM, NaHCO3 84 12mM, KHCO3 101.12mM, Taurine 125.1mM, Phenol red 376.4mM, Hepes 10mM) and then with digestion buffer (perfusion buffer containing 2.7 mg Liberase TM (Roche Life Sciences, Germany) for ~8 to 10mins at 37 °C. Mice were treated with anticoagulant (500U heparin i.p.) 30 min prior to heart extractions. After perfusion, the atria were removed and the heart was minced in the digestion buffer. Minced tissue was gently pipetted up and down on ice to release the cells, which were filtered through a 100μm pluristrainer mesh (pluriSelect Life Science, Leipzig, Germany) and then centrifuged for 5 min at 3,000 *g*. Cardiomyocytes and fibroblasts were separated using the pre-plating method and their purity was determined by qRT-PCR analysis. Cardiomyocytes isolated from two hearts were pooled and RNA was immediately extracted with Qiagen RNeasy kit (QIAGEN Inc., Germantown, Maryland).

**RNA sequencing and computational analysis**

Total RNA was purified from freshly isolated cardiomyocytes using RNeasy tissue mini-kit (Qiagen). RNA quality was analyzed with Agilent Bioanalyzer and quantified using a

Qubit fluorimeter at the Functional Genomics Core at the Roy J. Carver Biotechnology Center, UIUC. All RNA samples had at least A260nm/A280nm≥1.8, A260nm/A230nm≥1.4, r28S:16S≥1.5, and RNA integrated number (RIN)≥6.5. Hi-Seq4000 libraries were prepared, and 100-bp paired-end Illumina sequencing was performed on a HiSeq platform at the High Throughput Sequencing and Genotyping Unit, UIUC. Computational analysis of RNA-seq experiments for this study, and of previously published datasets **(Supplementary Table 2)** was performed as previously described[3]. Briefly, the sequencing reads were processed for quality and read length filters with Trimmomatic (version 0.38)[4]. RNA-seq reads were further aligned to the mouse (mm10) or human (hg38) genomes using STAR (version 2.4.2a)[5]. Mapping percentages and sample details are provided in **Supplementary Table 2**. Gene expression levels were determined as FPKM/TPM using count and differential expression values obtained from Cuffdiff[6]. Genes were considered as having significantly different expression following imposed cutoff clearance (FDR (q-value) <0.05, $\log_2$(fold change) >1). Differential splicing analysis was performed using rMATS (version 3.2.5), and significant events were identified with imposed cutoffs (FDR <0.10, junction read counts ≥20, PSI ≥20%)[7]. Motif analysis for differentially spliced exons was performed using rMAPS with default parameters, and by adding putative motifs as described previously[8,9]. Gene ontology analysis was performed with DAVID (version 6.8[10], and mapped using the 'Enrichment Maps' plugin in Cytoscape[11]. All expressed genes with TPM>1 served as background, and the biological function category was analyzed with three pathways (Biocarta, Kegg, and Panther). Functional clustering was performed and top clusters (*P* value <0.05) represented.

To identify co-regulated exons between RNA-seq samples from mouse and human experiments, the analysis was limited to mm10 annotated mouse cassette exons, which were converted to corresponding human exons in the hg38 annotation using UCSC liftover with minimum ratio of bases matching as 0.8. To identify $RBFOX2_{40}$-regulated exons in mouse cardiomyocytes that also contained $RBFOX2_{40}$ binding peaks, previously published iCLIP data for heavy molecular weight fraction (HMW) from mouse brains was used[12]. $RBFOX2_{40}$ binding clusters near these regulated exons were identified with the HOMER pipeline using its standard parameters[13]. To associate alternative exons with $RBFOX2_{40}$ binding, custom scripts were used to search for binding peaks present either in the exon or the surrounding upstream or downstream introns. To identify the crosslink sites for peaks associated with alternative splicing events, the CTK pipeline from CITS algorithm was used with default parameters[14,15]. After crosslink sites were identified, strand-specific sequences within a 70-nt window were obtained using getfasta script in the bedtools suite, which was followed by motif enrichment analysis using the MEME suite with standard parameters[16].

**Protein isolation and western blot analysis**

Protein lysates were prepared from indicated frozen heart tissues or freshly isolated cardiomyocytes by homogenization in bullet blender followed by sonication. Homogenization buffer (10 mM HEPES-KOH, pH 7.5, 0.32 M sucrose, 5 μM MG132, 5 mM EDTA, and Pierce proteinase inhibitor tablet (1 tablet/ 20 mL buffer volume) was used to isolate protein from heart tissues. Prior to sonication 20% Sodium Dodecyl sulfate (SDS) to a final concentration of 1% (v/v) was added. Protein concentrations were measured using Thermo scientific BCA assay kit. Approximately 50 µg of total protein sample was loaded onto a 10% SDS-PAGE gel, and then transferred onto a PVDF membrane overnight at 4$^{o}$C. Membranes were blocked using 5% milk powder (w/v) in TBST (Tris-buffered saline, 0.1% Tween 20) for 2 hours at room temperature (RT). Blots were incubated in primary antibodies at pre-determined concentrations for 2 hours. Blots were washed in TBST, and then incubated in HRP conjugated secondary antibodies for 1 hour at RT. Blots were developed using Clarity Western ECL kit (Bio-Rad). All antibodies used, and their respective dilutions are listed in **Supplementary Table 3**.

**Gene expression and splice isoform analysis**

Total RNA from mouse hearts and cardiomyocytes, were isolated using either RNeasy kit (QIAGEN Inc., Germantown, Maryland) or TRIzol reagent (Thermo Fisher Scientific, Waltham, Massachusetts). Upon DNase I treatment, 2 µg of RNA was reverse transcribed to cDNA using random hexamers and Maxima Reverse transcriptase kit (Thermo Fisher Scientific, Waltham, Massachusetts). The cDNA was diluted to a final concentration of 25 ng/µL and Real-time q-RTPCR and RT-PCR based alternative splicing assays were performed as described previously[17]. For splicing assays, the PCR products were analyzed by electrophoresis on a 6.5% polyacrylamide gel, stained with ethidium bromide and quantified using a Chemidoc XRS+ imaging system (Bio-Rad, Hercules, CA). PSI values for the variably spliced region were calculated with ImageLab software (BioRad) as [(exon inclusion band intensity) / (exon inclusion band intensity+exon exclusion band intensity) × 100]. All primers used for alternative splicing are listed in **Supplementary Table 1**. *SCN5A* exons 6A and 6B are of similar size (93-nts), therefore, PCR products were digested by *BstbI* enzyme before loading on 6.5% polyacrylamide gel, as previously described[18].

q-RTPCR was performed using the PerfeCTa SYBR® Green SuperMix (Quanta Biosciences, Beverly, Massachusetts) in an ABI QuantStudio3 (Thermo Fisher Scientific, Waltham, Massachusetts) with 10 min at 95 °C followed by 40 cycles of 15 s at 95 °C, 1:00min at 60°C using primers described in the **Supplementary Table 1**. *Gapdh* mRNA was used as a loading control and data were analyzed using the $2^{-\Delta\Delta Ct}$ method as previously described[19].

**miRNA profiling**

To quantify mature miRNA expression, we used TaqMan stem-loop RT-PCR MicroRNA Assays (Applied Biosystems). Briefly, 100 ng of total RNA from each sample was reverse-transcribed using specific miRNA primers from the TaqMan MicroRNA Assays and the TaqMan miRNA reverse transcription kit. Quantitative real-time PCR reactions were performed in triplicate on an ABI QuantStudio3 (Thermo Fisher Scientific, Waltham, Massachusetts) Real-Time PCR System using miRNA-specific TaqMan probe and ABI TaqMan Universal PCR Master Mix. An initial denaturation step of 10 min at 95°C was followed by 40 cycles of 95°C for 15 s and 60°C for 1 min. U6 snRNA was used to normalize the miRNA expression, and Fold change of miRNA expression was calculated as previously described[20].

**Cell culture and transfection**

HL-1 cardiomyocytes were cultured on gelatin (0.02%, w/v)/fibronectin (10µg/ml) coated plates with Claycomb Medium supplemented with 10% FBS (Sigma-Aldrich Corporation, St. Louis, Missouri) 2 mM L-glutamine, 0.1 mM Norepinephrine, 100 units/mL penicillin and 100 µg/mL streptomycin and were maintained at 37°C in 5% CO2 as previously descried[20]. After the cells reached 80-90% confluency and started beating, the cultures were split into new T75 flasks for maintenance or 12 well plates for further experiments. HL-1 cells were transiently transfected with DT0 and DT960 plasmids with Lipofectamine 3000 (Thermo Fisher Scientific, Waltham, Massachusetts) using manufacturer's protocol. To increase the transfection efficiency, we applied reverse transfection technique, where the plasmid DNA and transfection reagent complex is assembled in the tissue culture plate and then the cells are seeded into the wells. Forty-eight hours later cells were harvested to isolate total RNA and protein. *Celf1* siRNA delivery was performed 12 h after DT0 or DT960 plasmid transfections, using Lipofectamine RNAi-Max (Thermo Fisher Scientific, Waltham, Massachusetts) according to manufacturer's recommendations. Final concentration of siRNAs was 10nM. The sequences/reference numbers of siRNAs used are listed in **Supplementary Table 1**. To induce *Rbfox2* 43-nt skipping, an antisense morpholino oligomer (*Rbfox2*-43 ASO) sequence was used: 5'-CACTGGCACTCCTAC CTGAGGTATT-3', which binds to the 5'ss of *Rbfox2* exon 43 pre-mRNA sequence. As a control, a non-target standard control sequence was used:5'-CCTCTTACCTCAGTTACAATTTATA-3' (Gene Tools, LLC, Philomath, Oregon, USA).

Morpholinos were delivered at 10uM final concentration using Endo-porter reagent in HL-1 cells for 48h and cells were harvested to prepare total RNA and protein lysates.

**Luciferase activity assays**

The 3'-UTR of mouse *Rbfox2* was cloned at the *Not1* and *Xho1* site of the psiCHECK$^{TM}$-2 plasmid (Promega), downstream of *Renilla* Luciferase (hRLuc). The primers used to amplify the *Rbfox2* 3'-UTR are listed in **Supplementary Table 1**. To generate the indicated luciferase constructs, In-Fusion HD cloning system (Clontech) was used. Next, we introduced mutations in *Let-7*, *miR-9* and *miR-135a* binding sites within the *Rbfox2* 3'-UTR construct by using the QuikChangeII Mutagenesis Kit (Stratagene, La Jolla, CA, USA). A combined *Let-7+miR-9+miR-135a* mutant construct was also generated in which all three miRNAs binding sites were mutated simultaneously. The primers used for incorporating mutations were designed using Quick-change primer design software and are listed in **Supplementary Table 1**. HEK293T cells were seeded on twelve-well plates at approximately 70–80% confluence and after 24h, 1µg of luciferase expression construct and 20nM of miRIDIAN microRNA Mimics (Dharmacon, USA) were co-transfected with *Trans*IT-X2 (Mirus Bio LLC, USA). A Non-targeting mimic was used as control (miRIDIAN microRNA Mimic Negative Controls, CN-001000-01-05, Dharmacon, USA). After 48h from transfection, the cells were lysed and assayed with Dual Luciferase Assay (Promega) according to the manufacturer's instructions. Wildtype and mutant *Rbfox2* 3'-UTR constructs were tested in three independent experiments. Renilla luciferase activity was normalized with the Firefly luciferase activity levels and expressed as relative luciferase units (RLU). For the rescue experiments, HL-1 cells were reverse-transfected with DT0 or DT960 plasmids. Twelve hours later, *Rbfox2* 3'-UTR reporter plasmid was co-transfected with a cocktail of 10nM *Let-7g+miR-9+miR-135a* miRNA mimics with *Trans*IT-X2 (Mirus Bio LLC, USA). Cells were incubated for 48h, collected for RNA and protein extractions, and luciferase activity assays.

**Adenovirus production**

cDNAs encoding FLAG-tagged human CELF1 were sub-cloned into the p-Adeno-X-ZsGreen1 vector (Clonetech, 632267) using the In-Fusion kit (Clonetech, 639646) as per the manufacturer's instructions. High-titer adenoviruses were generated and purified as mentioned before[8]. To determine the effect of CELF1 overexpression, six-wells containing ~70% confluent HL-1 cells were infected with $2.0 \times 10^9$ o.p.u. (optical particle units) of the CELF1 or GFP adenovirus for 48h and cells were harvested to extract RNA and protein for further analysis.

**Homology modeling**

The cryo-EM structures of eukaryotic sodium channels $Na_v1.4$ (pdb:6agf) and $Na_vPas$ (pdb:6a95) were used as templates for modeling a human sodium channel $Na_v1.5$ (SCN5A, UniProtKB: Q14524). The sequence segments corresponding to the domain I-II linker and domain II-III linker were truncated from the full human SCN5A protein sequence to allow better alignment with the templates. Multi-sequence alignment between the templates and target sequence was performed using Clustal Omega[21] and the homology modeling was performed using MODELLER v9.21[22]. 50 models were generated for both the adult and fetal isoforms of SCN5A and the model with the best objective function score was selected for each isoform to perform molecular dynamics (MD) simulations.

**MD simulations**

The selected homology models for both isoforms were embedded in a POPC lipid bilayer and solvated with 0.15 M of NaCl and TIP3P water[23] using CHARMM-GUI Membrane Builder[24]. The dimension for both systems is 140 Å×140 Å×145 Å. Both systems underwent 1 ns NPT initial equilibration with the standard protocol described in the CHARMM-GUI Membrane Builder, which involves gradually releasing positional and dihedral restraints on the protein and lipid molecules. Thereafter, 10 ns of NPT equilibration with dihedral restraints (k=100 kcal/mol/rad$^2$) on the protein secondary structure were performed. The last frame of each system was then used for three independent (different initial velocities) NPT simulations. To remove any initial structural bias from the initial model, each independent trajectory was further equilibrated without restraint for 20 ns, followed by 100 ns production run. All simulations were carried out with NAMD 2.13[25], using CHARMM36m protein[26] and CHARMM36 lipid[27] parameters. SHAKE algorithm[28] was employed to constrain hydrogen bond lengths to allow 2 fs time steps for the integrator. Constant temperature of 310 K was maintained by Langevin thermostat[29] with a damping coefficient of 1 ps$^{-1}$. Nosé-Hoover Langevin piston[30] with a period of 200 ps and decay time of 50 ps was employed to maintain constant pressure at 1 atm. Periodic boundary condition and a nonbonded cutoff of 12 Å (with 10 Å switching distance and vdW force switching) were used. Long-range electrostatics were calculated using the particle mesh Ewald method[31] with 1 Å grid spacing.

**Statistics**

All experiments have at least three independent biological repeats. Differences between groups were examined for statistical significance using Student's t-test with Welch's correction (for two groups), or one-way ANOVA plus Dunnett's post-hoc test (for more than two groups) using the GraphPad Prism 7 Software. Results were expressed as mean ± s.d., unless otherwise specified. *$p<0.05$, **$p<0.001$, were considered statistically significant.
